# Intraneuronal chloride accumulation via NKCC1 is not essential for hippocampal network development *in vivo*

**DOI:** 10.1101/2020.07.13.200014

**Authors:** Jürgen Graf, Chuanqiang Zhang, Stephan Lawrence Marguet, Tanja Herrmann, Tom Flossmann, Robin Hinsch, Vahid Rahmati, Madlen Guenther, Christiane Frahm, Anja Urbach, Ricardo Melo Neves, Otto W. Witte, Stefan J. Kiebel, Dirk Isbrandt, Christian A. Hübner, Knut Holthoff, Knut Kirmse

## Abstract

NKCC1 is the primary transporter mediating chloride uptake in immature principal neurons, but its role in the development of *in vivo* network dynamics and cognitive abilities remains unknown. Here, we address the function of NKCC1 in developing mice using electrophysiological, optical and behavioral approaches. We report that NKCC1 deletion from telencephalic glutamatergic neurons decreases *in-vitro* excitatory GABA actions and impairs neuronal synchrony in neonatal hippocampal brain slices. *In vivo*, it has a minor impact on correlated spontaneous activity in the hippocampus and does not affect network activity in the intact visual cortex. Moreover, long-term effects of the developmental NKCC1 deletion on synaptic maturation, network dynamics and behavioral performance are subtle. Our data reveal a neural network function of depolarizing GABA in the hippocampus *in vivo*, but challenge the hypothesis that NKCC1 is essential for major aspects of hippocampal development.

## Introduction

Intracellular chloride concentration ([Cl^−^]_i_) is a major determinant of neuronal excitability, as GABAergic inhibition is primarily mediated by chloride-permeable ionotropic receptors (Bormann et al., 1987). In the mature brain, [Cl^−^]_i_ is maintained at low levels by secondary active chloride extrusion, which renders GABA hyperpolarizing (Rivera et al., 1999) and counteracts activity-dependent chloride loads (Doyon et al., 2016). Effective GABAergic inhibition in the adult brain is crucial not only for preventing runaway excitation of recurrently connected glutamatergic cells (Krnjevic et al., 1963) but also for entraining neuronal assemblies into oscillatory rhythms underlying cognitive processing (Allen et al., 2015). However, it has long been known that the capacity of chloride extrusion is low during early brain development (Luhmann et al., 1991; Spoljaric et al., 2017). In addition, immature neurons are equipped with chloride uptake mechanisms, in particular with the electroneutral Na^+^/K^+^/2Cl^−^ cotransporter NKCC1 (Dzhala et al., 2005; Pfeffer et al., 2009; Sipilä et al., 2009; Wang et al., 2008; Yamada et al., 2004). NKCC1 contributes to the maintenance of high [Cl^−^]_i_ in the intact developing brain (Sulis Sato et al., 2017), thus favoring depolarizing responses to GABA_A_ receptor (GABA_A_R) activation *in vivo* (Kirmse et al., 2015; van Rheede et al., 2015).

When GABA acts as a mainly depolarizing neurotransmitter, neural circuits generate burst-like spontaneous activity (Ackman et al., 2012; Arichi et al., 2017; Khazipov et al., 2004; Kummer et al., 2016; Leinekugel et al., 2002), which is thought to be crucial for their developmental refinement (Blanquie et al., 2017; Oh et al., 2016; Winnubst et al., 2015; Zhang et al., 2011). A large body of evidence from acute brain slices indicates that cortical GABAergic interneurons promote neuronal synchrony in an NKCC1-dependent manner (Dzhala et al., 2005; Flossmann et al., 2019; Pfeffer et al., 2009; Rheims et al., 2008; Valeeva et al., 2016; Wester et al., 2016). However, the *in vivo* developmental functions of NKCC1 are far from understood (Han et al., 2015; Kirmse et al., 2018). One fundamental question in this context is to what extent NKCC1-dependent GABAergic depolarization instructs the generation of correlated spontaneous activity in the intact developing brain. In the rodent neocortex *in vivo*, GABAergic transmission imposes spatiotemporal inhibition on spontaneous and sensory-evoked activity already in the neonatal period (Che et al., 2018; Colonnese et al., 2010; Kirmse et al., 2015; Valeeva et al., 2016). Whether a similar situation applies to other brain regions is unknown, as two recent studies employing chemo- and optogenetic techniques for manipulating GABA release from hippocampal interneurons yielded opposing results (Murata et al., 2020; Valeeva et al., 2016). Experimental manipulations of the chloride driving force are potentially suited to resolve these divergent findings, but available pharmacological (Löscher et al., 2013; Marguet et al., 2015; Sipilä et al., 2006) or conventional knockout (Delpire et al., 1999; Pfeffer et al., 2009; Sipilä et al., 2009) strategies suffer from unspecific effects that complicate interpretations.

Here, we overcome this limitation by selectively deleting *Slc12a2* (encoding NKCC1) from telencephalic glutamatergic neurons. Using optical and electrophysiological recordings, we show that chloride uptake via NKCC1 strongly promotes synchronized network activity in acute hippocampal slices *in vitro*, but has weak and event type-dependent effects on spontaneous network activity in the neonatal CA1 *in vivo*. The long-term loss of the cotransporter leads to subtle changes of network dynamics in the adult hippocampus, leaving synaptic development unperturbed and hippocampus-dependent behavioral performance intact. Our data suggest that NKCC1-dependent chloride uptake is largely dispensable for several key aspects of hippocampal development *in vivo*.

## Results

### Behavioral performance of mice in the conditional absence of NKCC1 from telencephalic glutamatergic cells

To investigate the role of depolarizing GABAergic transmission in telencephalic glutamatergic neurons, we conditionally deleted *Slc12a2* in *Emx1*-lineage cells of embryonic mice using a transgenic Cre-loxP-based approach (Figure 1A). A comparison to *NKCC1^flox/flox^* (WT) littermates showed that Cre-dependent recombination had been widespread in the hippocampus (Figure 1B) and led to a profound reduction in total hippocampal *Slc12a2* mRNA levels in *Emx1^IREScre^:NKCC1^flox/flox^* (KO^Emx1^) mice as early as at postnatal days (P) 2–3 (Figure 1C and Table S1). Fate mapping of *Emx1*-lineage cells in hippocampal areas CA3 and CA1 at P54–63 confirmed that Cre expression was restricted to glutamatergic neurons (GAD67^−^/NeuN^+^), whereas cells expressing either a pan-GABAergic (GAD67) or a subtype-selective GABAergic interneuron (parvalbumin or somatostatin) marker were Cre-reporter negative (Figure S1A–J). WT and KO^Emx1^ mice were born at Mendelian ratios (P = 0.41, χ² = 0.69, n = 71 mice, chi-squared-test), displayed a normal postnatal gain in body weight (Figure S1K), similar brain growth (measured by lambda-bregma distance, see #14 in Table S1), and showed unperturbed developmental reflexes, including cliff avoidance behavior and righting reflex (Figures S1L and S1M). Next, we subjected adult mice to a battery of behavioral tests to assess whether the conditional loss of NKCC1 would result in an overt phenotype. Unlike conventional NKCC1 knockout mice (Antoine et al., 2013), KO^Emx1^ mice exhibited unaltered locomotor activity and exploratory behavior as assessed in the open field (Figures 1D and 1E). Spontaneous alternations in a Y-maze, which served as a measure of spatial working memory, were unaffected (Figures 1F and 1G). Spatial learning and memory as evaluated in a Morris water maze (Figures 1H–1K), as well as cue- and context-specific fear memory (Figures 1L–1N), were not significantly altered by the chronic loss of NKCC1, either. In sum, KO^Emx1^ mice lack the severe phenotype of NKCC1 null mice, which is due to NKCC1 loss from tissues other than the brain (Delpire et al., 1999). In addition, the phenotype resulting from the conditional deletion of NKCC1 from telencephalic principal neurons is compatible with a normal development of several hippocampus-dependent and -independent behaviors.

**Figure 1.**
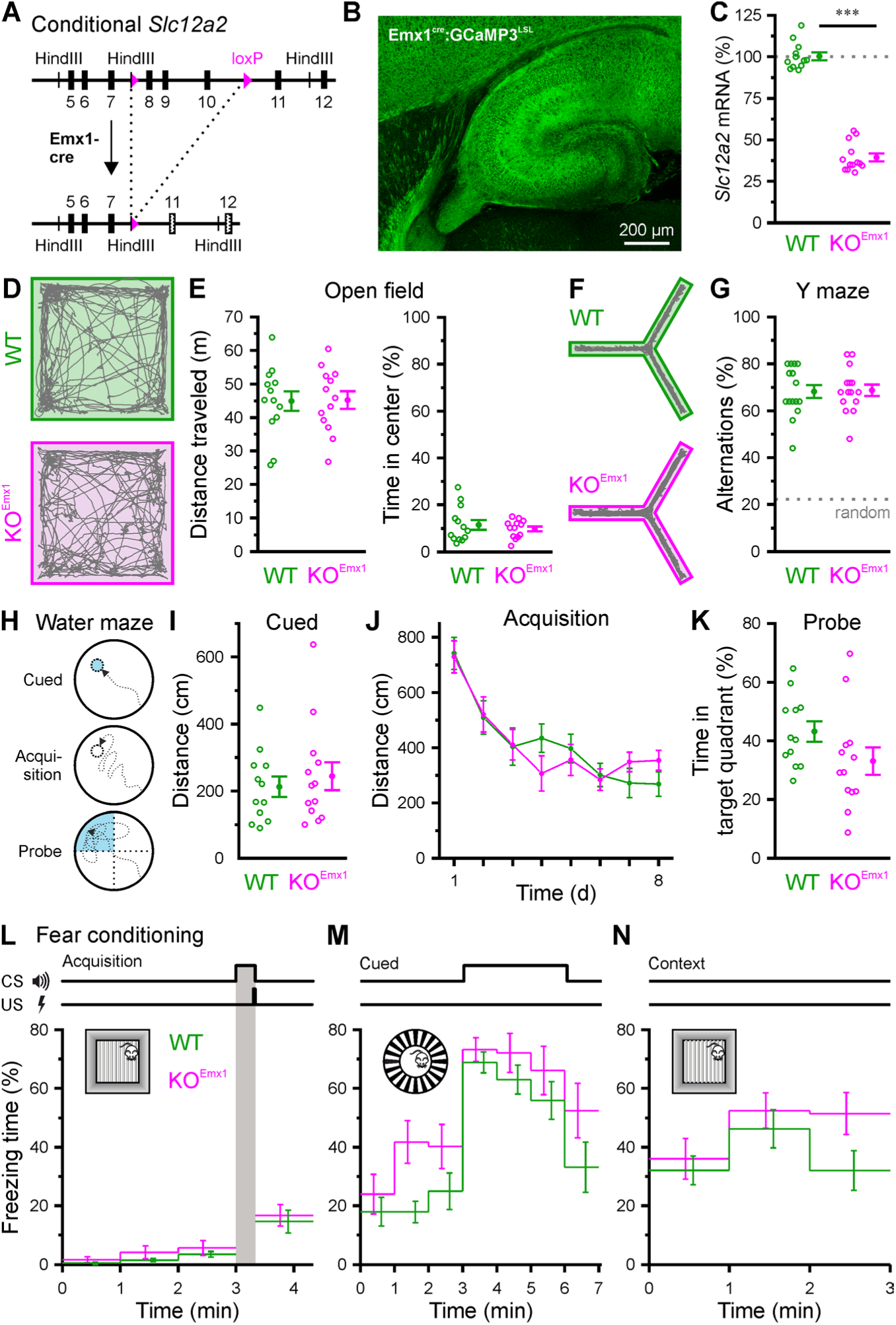
Behavioral performance of mice in the absence of NKCC1 from telencephalic glutamatergic cells. (A) In the targeted *Slc12a2* locus, exons 8–10 are flanked by loxP sites. Cre-dependent recombination causes a frameshift (dotted exons) and introduces a stop codon in exon 12. (B) Confocal image demonstrating Cre-reporter expression (GCaMP3) in a fixed horizontal brain slice obtained from an *Emx1^IREScre^:Ai38* mouse at P4. (C) Hippocampal *Slc12a2* mRNA levels compared to the geometric mean of WT, normalized to *Gapdh* and *Hmbs*. (D) Sample trajectories of mice in the open field. (E) Total distance covered and relative time the animals spent in the center. (F) Sample trajectories of mice in the Y maze. (G) Spontaneous alternations (dotted line indicates chance level). (H) Experimental design of the Morris water maze. (I, J) Distance to platform for cued trial (visible platform) and acquisition (hidden platform). (K) Time in target quadrant during probe trial (no platform). (L–N) Freezing time during fear conditioning (L), and during re-exposure to the cue (M) and context (N). (C–K) Each open circle represents a single animal. (E–N) Data are presented as mean ± SEM. CS – conditioned stimulus, US – unconditioned stimulus, ^***^ P < 0.001. See also Figure S1 and Table S1.

### Loss of NKCC1 attenuates the depolarizing action of GABA

We next investigated how the loss of NKCC1 alters the cellular actions of GABA in the neonatal hippocampus. To this end, we first used low-gramicidin perforated-patch current-clamp recordings that allow monitoring the membrane potential (V_m_), while minimizing the risk of unintended membrane breakthrough (Zhang et al., 2019; Zhu et al., 2008). Measurements were performed in acute slices (P3–4) in the presence of antagonists of ionotropic glutamate receptors (10 µM DNQX, 50 µM APV) and voltage-gated Na^+^ (0.5 µM TTX) and Ca^2+^ (100 µM CdCl_2_) channels so as to block recurrent excitation and amplification of voltage changes by intrinsic conductances. Resting membrane potential of CA3 pyramidal cells (PCs) was similar in WT and KO^Emx1^ mice (Figures 2A and 2B, Table S2). However, NKCC1 deletion resulted in a significantly reduced peak membrane potential (V_peak_) induced by a saturating puff of the GABA_A_ receptor (GABA_A_R) agonist isoguvacine (100 µM, 2 s) by almost 10 mV (WT: –50.9 ± 1.9 mV, n = 15, KO^Emx1^: –58.6 ± 2.8 mV, n = 10, P = 0.028, two-sample t-test; Figure 2B). Because V_peak_ will approximate the reversal potential of GABA_A_R-mediated currents (E_GABA_) under these conditions, our data imply lower [Cl^−^]_i_ in cells obtained from NKCC1 KO^Emx1^ mice. Isoguvacine-induced V_m_ changes in individual cells were consistent across successive trials (Figure 2C). In line with this observation, amplitudes of somatic Ca^2+^ transients (CaTs) evoked by bath application of GABA (100 µM, 1 min) in the continuous presence of TTX were attenuated in putative PCs of KO^Emx1^ mice (P2–4), whereas glutamate-induced CaTs were unaffected (Figure S2). Furthermore, cell-attached voltage-clamp recordings in the presence of ionotropic glutamate receptor blockers (10 µM DNQX, 50 µM APV) revealed that puff application of isoguvacine (100 µM, 100 ms) transiently increased action potential (AP) frequency in WT cells, but this effect was largely absent from PCs of KO^Emx1^ mice at P3–4 (Figures 2D–2F). We conclude that NKCC1 deletion attenuates GABA_A_R-mediated depolarization and virtually abolishes excitatory effects of GABAergic transmission *in vitro*.

**Figure 2.**
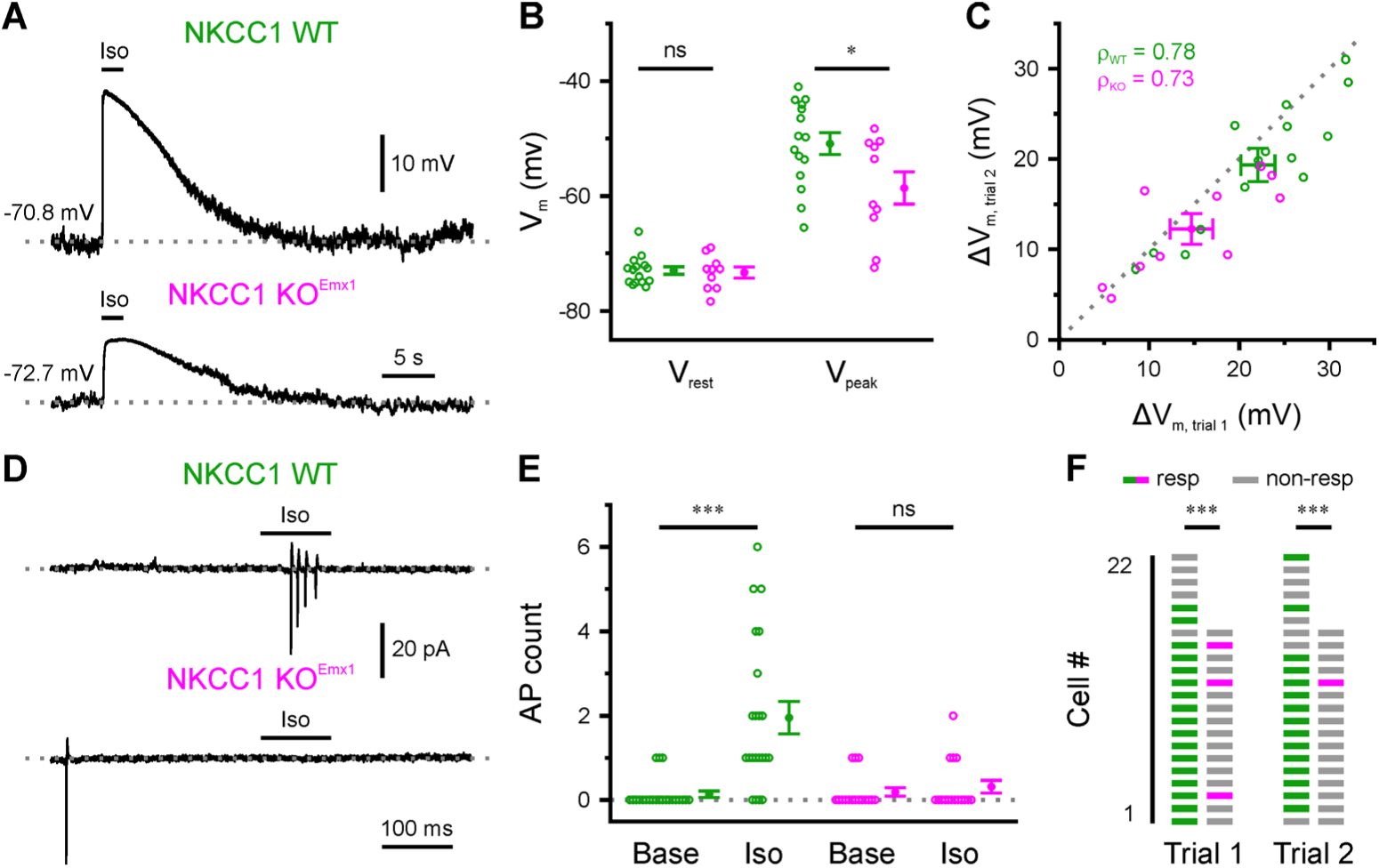
Loss of NKCC1 attenuates the depolarizing action of GABA. (A) Sample gramicidin perforated-patch current-clamp recordings in response to puff application of isoguvacine (Iso). (B) Quantification of resting (V_rest_) and peak isoguvacine-induced (V_peak_) membrane potential. (C) Correlation of isoguvacine-induced membrane potential changes (ΔV_m_) in two successive trials. ρ – Spearman’s rank correlation coefficient. (D) Sample cell-attached recordings in response to puff-application of isoguvacine. (E) Number of action currents detected in 300-ms intervals immediately before (Base) and after (Iso) puff onset. (F) The fraction of responsive cells (resp) was significantly smaller for cells from KO^Emx1^ mice. Each open circle represents a single cell. Data are presented as mean ± SEM. ns – not significant, ^*^ P < 0.05, ^***^ P < 0.001. See also Figure S2 and Table S2.

### PC-specific NKCC1 deletion impairs spontaneous hippocampal network activity in vitro

Based on previous experimental and theoretical results (Flossmann et al., 2019; Pfeffer et al., 2009; Sipilä et al., 2006), we hypothesized that a PC-specific attenuation of GABAergic depolarization would impair synchronized network activity. We tested this prediction in the CA3 area of acute hippocampal slices (P1–4) using whole-cell voltage-clamp recordings of spontaneous GABAergic postsynaptic currents (sGPSCs) isolated by their reversal potential (Figure 3A). We found that sGPSC frequencies did not significantly differ between genotypes (Figure 3B and Table S3). In WT slices, sGPSCs typically occurred in distinct bursts [also known as giant depolarizing potentials (GDPs) (Ben-Ari et al., 1989)] (Figure 3A). However, this temporal structure was largely absent in cells recorded from KO^Emx1^ mice (Figure 3A). To quantify this effect, a frequency-dependent metric referred to as burst index (see Methods) was computed for measured data and compared to a Poisson point process in which sGPSCs occur randomly. Irrespective of the average sGPSC frequency, burst indices of WT, but not of KO^Emx1^, cells were consistently higher than those expected for a random (Poisson) case (Figure 3C). When controlling for the effect of sGPSC frequency, we found that burst indices were significantly reduced by the conditional loss of NKCC1 (main effect genotype: P < 1.2 × 10^−4^, F = 26.3, η_p_² = 0.64, df = 1, analysis of covariance; Figure 3D), suggesting that neuronal synchrony was profoundly impaired in KO^Emx1^ mice. This conclusion was confirmed by single-cell confocal Ca^2+^ imaging using the synthetic indicator OGB1, which revealed a dramatic reduction in GDP frequency in CA3 of KO^Emx1^ mice at P1–4 (Figure S3). Of note, the effects of a conditional loss of NKCC1 were region-dependent because the kinetics of single-cell recruitment in CA1 during GDPs was significantly slowed down in slices from KO^Emx1^ mice, while the GDP frequency was similar in both genotypes (Figure S3).

**Figure 3.**
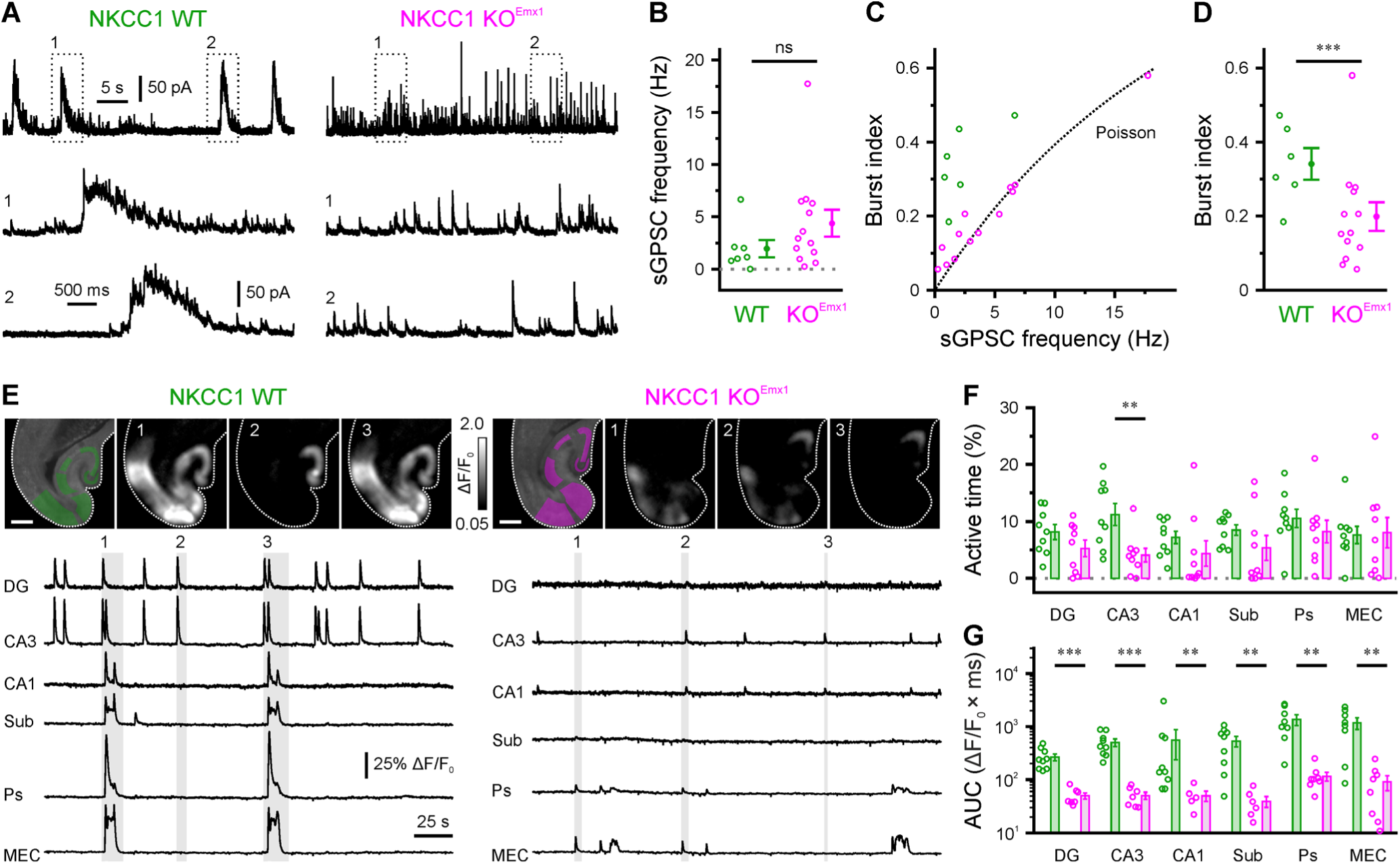
PC-specific NKCC1 deletion impairs spontaneous hippocampal network activity *in vitro*. (A) Sample whole-cell recordings of sGPSCs isolated by their reversal potential. (B) Quantification of sGPSC frequency. (C) Burst index as a function of sGPSC frequency. The dotted line represents the expected value for stochastic event trains following a Poisson point process. (D) Quantification of burst indices. (E) GCaMP-based confocal imaging of spontaneous network activity in cortico-hippocampal slices. *Top*: GCaMP3 fluorescence overlaid with regions of interest used for analysis (left) and sample ΔF/F_0_ images for time periods indicated below (scale bars, 500 µm). *Bottom*: ΔF/F_0_ sample traces. (F–G) Quantification of total active time (F) and mean area under the curve per cluster event (AUC, G). Open circles represent single cells (B–D) or slices (F–G). Data are presented as mean ± SEM. ns – not significant, ^*^ P < 0.05, ^**^ P < 0.01, ^***^ P < 0.001. See also Figure S3 and Table S3.

We next sought to extend these findings by performing independent large-scale confocal Ca^2+^ imaging experiments using a conditional transgenic expression of the genetically encoded indicator GCaMP3 in *Emx1*-lineage cells. At P1–4, large-amplitude network events were detected in slices obtained from WT (*Emx1^IREScre^:NKCC1^wt/wt^:GCaMP3^LSL^*) mice, which often occurred almost simultaneously in multiple hippocampal subfields and frequently involved both hippocampal and adjacent cortical areas (Figure 3E). While such activity was also present in slices from KO^Emx1^ (*Emx1^IREScre^:NKCC1^flox/flox^:GCaMP3^LSL^*) mice, the area under the curve of detected network events was substantially reduced in both the hippocampus and adjacent cortical regions (Figures 3E–3G). In addition, a reduction in the average active time was found in CA3 (Figure 3F). Collectively, several lines of evidence show that PC-specific NKCC1 deletion leads to a profound attenuation of neuronal synchrony *in vitro*.

### NKCC1 effects on hippocampal network dynamics are not accounted for by alterations in intrinsic excitability or basic synaptic properties

The question how GABAergic transmission instructs proper neuronal development remains largely unanswered (Kirmse et al., 2018). Some previous data indicate that the absence of NKCC1 could interfere with the normal maturation of synaptic (Wang et al., 2008) and/or intrinsic (Sipilä et al., 2009) conductances. We therefore set out to examine whether altered network dynamics observed in NKCC1 KO^Emx1^ mice result from an acute shift in E_GABA_ as reported above or, alternatively, from developmental changes following NKCC1 disruption in embryonic life. As an overall measure of intrinsic excitability, we first assessed spontaneous AP firing in the presence of ionotropic glutamate and GABA receptor antagonists (10 µM DNQX, 50 µM APV, 10 µM bicuculline methiodide [BMI]) by means of cell-attached voltage-clamp recordings from CA3 PCs at P3–4. In contrast to results obtained from an NKCC1 null line (Sipilä et al., 2009), AP frequencies were unaffected by the conditional loss of NKCC1 (WT: 0.31 ± 0.09 Hz, n = 8, KO^Emx1^: 0.29 ± 0.11 Hz, n = 11, P = 0.39, exact Mann-Whitney U-test; Figures 4A and 4B, Table S4). Moreover, NKCC1 disruption did not significantly alter neuronal input-output relationships as revealed by whole-cell recordings of AP firing in response to somatic current injections of variable amplitudes (Figures 4C and 4D). In agreement with this finding, neither passive membrane properties (see #10–12 in Table S4) nor AP threshold (Figure 4E) of CA3 PCs were modified by the loss of NKCC1.

**Figure 4.**
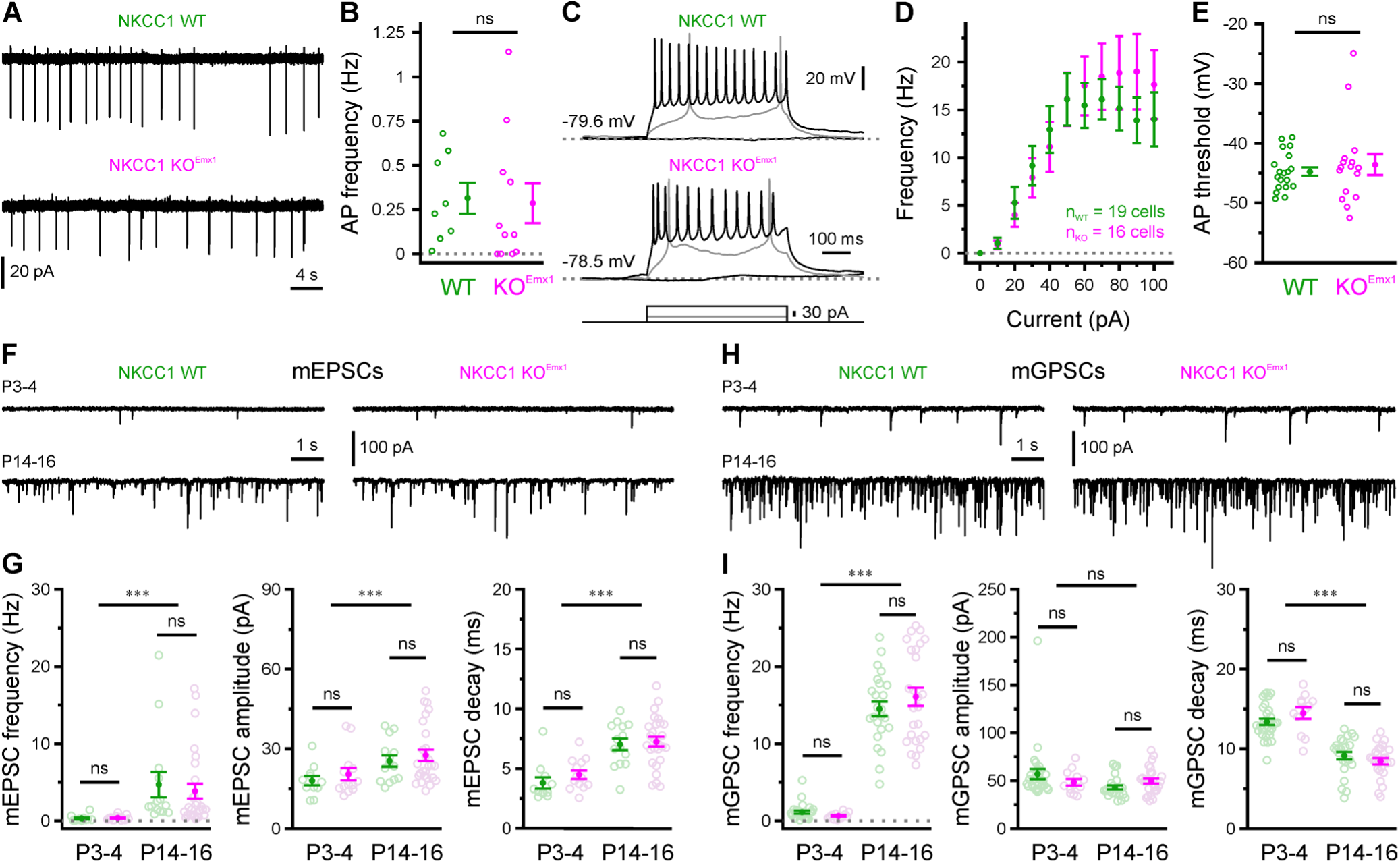
NKCC1 effects on hippocampal network dynamics are not accounted for by alterations in intrinsic excitability or basic synaptic properties. (A) Sample cell-attached recordings of spontaneous AP firing in the presence of ionotropic glutamate and GABA receptor antagonists. (B) Mean spontaneous AP frequency. (C) Sample whole-cell current-clamp recordings in response to current injections. (D–E) Current-frequency relationship (D) and AP threshold (E) are not altered in NKCC1 KO^Emx1^ mice. (F) Sample voltage-clamp measurements demonstrating AMPAR-dependent mEPSCs at P3–4 and P14–16. (G) Mean mEPSC frequency, median mEPSC amplitude and mean mEPSC decay time constant. (H) Sample voltage-clamp measurements demonstrating mGPSCs at P3–4 and P14–16. (I) Mean mGPSC frequency, median mGPSC amplitude and mean mGPSC decay time constant. Each open circle represents a single cell. Data are presented as mean ± SEM. ns – not significant, ^***^ P < 0.001. See also Table S4.

We next investigated whether the basic properties of glutamatergic and/or GABAergic synapses were altered upon PC-specific NKCC1 disruption. Whole-cell voltage-clamp recordings from CA3 PCs were performed at P3–4 and P14–16, because this time period is characterized by intense synaptogenesis and subunit reorganization of postsynaptic receptors. AMPA receptor-dependent miniature PSCs (mEPSCs) displayed a profound developmental increase in frequency, accompanied by a moderate increase in (quantal) amplitude and a deceleration in decay kinetics (Figure 4G) – in line with published data (Stubblefield et al., 2010). Of note, these maturational alterations were independent of the genotype, since no significant interaction (age × genotype) was found for any of the parameters examined (at least P > 0.6, two-way ANOVAs, Table S4). GABA_A_R-mediated miniature postsynaptic currents (mGPSCs) also underwent a substantial developmental increase in frequency (Figures 4H and 4I, left), along with an acceleration in their decay kinetics (Figure 4I, right) (Cohen et al., 2000), but no change in quantal amplitude (Figure 4I, middle). These developmental trajectories were indistinguishable between WT and KO^Emx1^ mice (two-way ANOVAs, Table S4). In summary, both intrinsic excitability and basic synaptic properties were unaffected by the conditional disruption of NKCC1 and, thus, are unlikely to account for the alteration of *in vitro* hippocampal network dynamics observed in NKCC1 KO^Emx1^ mice.

### Hippocampal sharp waves in vivo persist in the conditional absence of NKCC1

Next, we investigated the effect of the loss of NKCC1 in principal neurons on *in vivo*, awake hippocampal network and CA1 multi-unit activity (MUA) at P4. We recorded local field potentials (LFPs) along the CA1-dentate gyrus axis using linear 32-site silicon probes in awake head-fixed neonates. We observed similar activity patterns in P4 hippocampus of both WT and NKCC1 KO^Emx1^ mice: sharp waves (SPWs) with a phase reversal in the CA1 pyramidal layer and short-lasting oscillations in the 10–30 Hz (beta) frequency range (hippocampal network oscillations, HNOs; Figures 5A and 5B) as previously described (Fazeli et al., 2017; Marguet et al., 2015). Depth profiles of SPWs were comparable to those described in rat pups (Valeeva et al., 2019) (’rad/lm SPW’ in Figure 5C). In addition, we found inverted SPWs with their maximum negativity and a corresponding current sink in *stratum oriens* (s.o., ’oriens SPW’ in Figure 5C). Both SPW types elicited distinct MUA patterns in the CA1 pyramidal layer that were similar for both genotypes. None of these SPW events’ properties such as occurrence frequency and amplitudes or event durations differed between WT and KO^Emx1^ animals (Figure 5D and Table S5). Likewise, SPW properties did not differ between genotypes in P7 or P12 neonates (Figure S4). Furthermore, neither the rate nor the length of HNOs differed significantly between the genotypes at P4 (Figure 5D).

**Figure 5.**
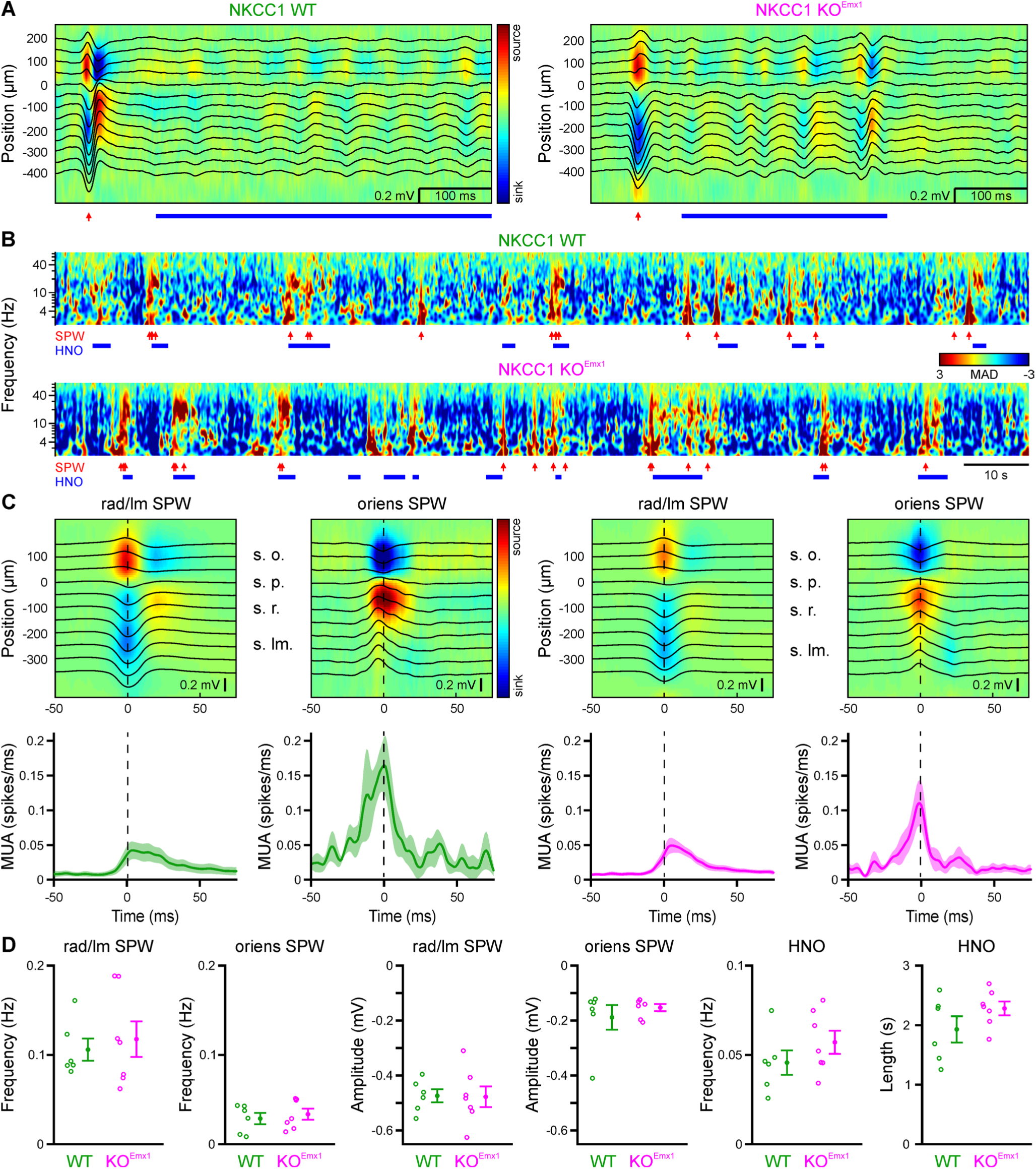
Hippocampal sharp waves *in vivo* at P4 persist in the conditional absence of NKCC1. (A) Example SPW (red arrow) and beta-frequency (∼10–30 Hz range) hippocampal network oscillation (HNO, blue line) recorded by linear silicon probes with 50 µm contact spacing, spanning the hippocampal layers. LFP traces (black) are superimposed over the current source density (CSD). 0 µm represents the field reversal of s.p. (B) Example 150 s wavelet spectrograms of CSD in s.l.m. in median absolute deviation (MAD) units per frequency. Events are marked as in (A) and (B). (C) Median LFP and CSD depicted from SPWs with sinks either below (’rad/lm’, left) or above (’oriens’, right) s.p. Left two panels are from a WT, and right two are from a KO^Emx1^ mouse. Lower panels display the SPW trough-aligned average multi-unit activity (MUA) for all sets with at least 10 events of that type present. (D) Summary statistics for the frequency and amplitude of rad/lm and oriens SPWs across all NKCC1 WT and KO^Emx1^ mice recorded. For HNOs the occurrence frequency as well as average duration is shown. Each open circle represents a single animal. Data are presented as mean ± SEM. See also Figure S4 and Table S5.

### NKCC1 promotes correlated network activity in an event type- and region-specific manner in vivo

As LFPs mainly result from postsynaptic current flow (i.e., neuronal input), we further characterized *in vivo* network dynamics using wide-field epifluorescence Ca^2+^ imaging, which primarily reflects AP firing (i.e., neuronal output). To gain optical access to CA1 *in vivo*, a hippocampal window preparation (Mizrahi et al., 2004) was adapted for use in neonatal mice (P3–4; Figure 6A). Control experiments based on LFP recordings, using a tungsten electrode positioned in *stratum radiatum* (s.r.), confirmed that SPW frequencies measured either through the intact cortex or after focal cortex aspiration did not significantly differ from each other (P = 0.91, exact Mann-Whitney U-test; Figures S5A–S5C and Table S6). Ca^2+^ imaging was then performed in mice conditionally expressing GCaMP3 in *Emx1*-lineage cells (Figure 6B). Under N_2_O anesthesia, spontaneous network activity occurred in the form of spatiotemporal clusters of CaTs, frequently recruiting large parts of the field of view (∼0.6 mm²), in all WT and KO^Emx1^ animals examined (n = 22; Figures 6B–6E). Simultaneous LFP recordings revealed that approximately two-thirds of Ca^2+^ clusters were temporally unrelated to SPWs (–SPW; Figures 6C–6E and 6G; see Methods) in both genotypes. –SPW Ca^2+^ clusters lacked a consistent electrophysiological signature (Figures 6C and 6D), suggesting that LFP measurements are biased towards SPW-related (+SPW) network events in CA1. Importantly, both +SPW and –SPW Ca^2+^ clusters were entirely AP-dependent (Figures S5D and S5E).

**Figure 6.**
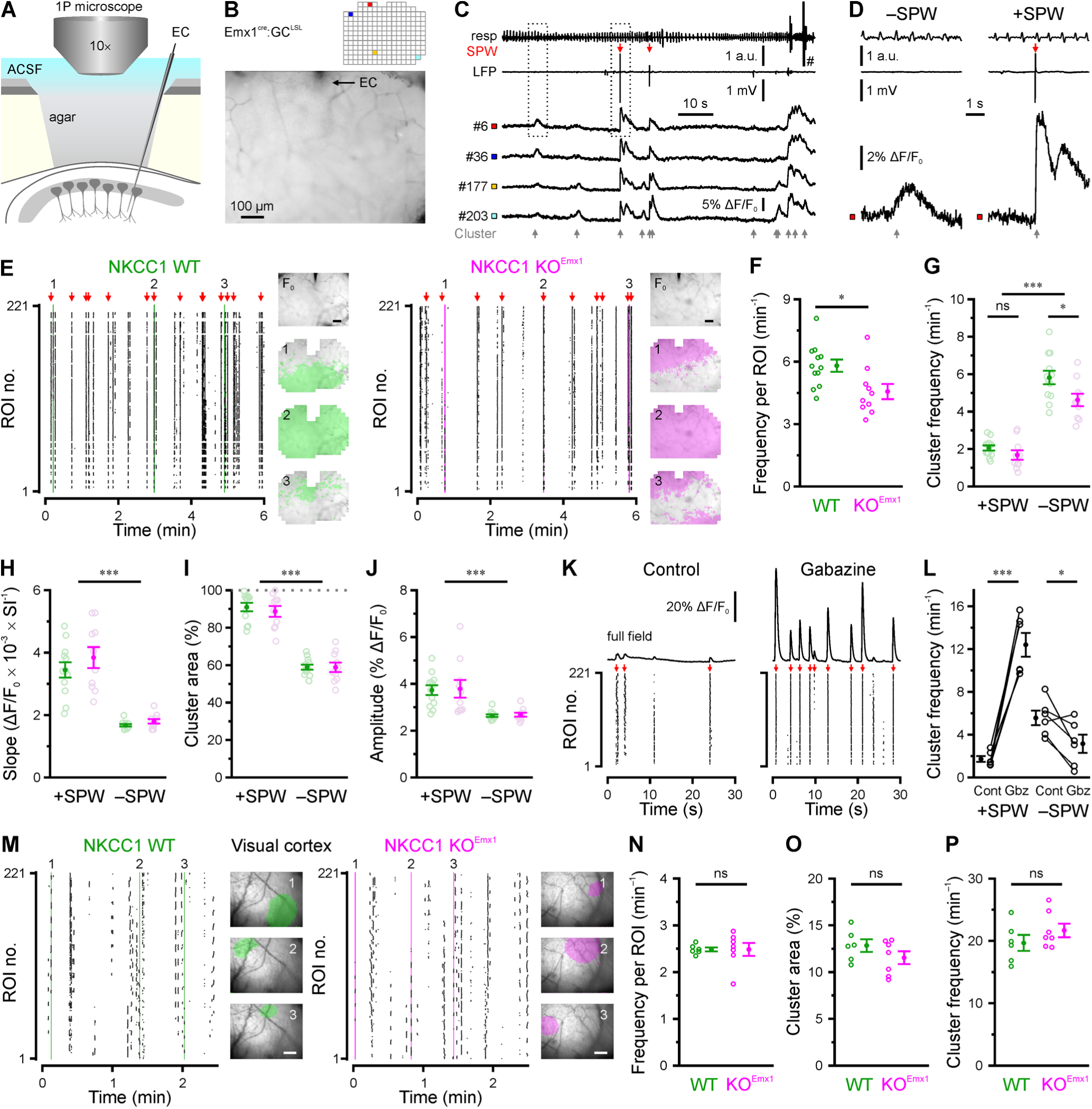
NKCC1 promotes correlated network activity in an event type- and region-specific manner *in vivo*. (A) Experimental set-up combining wide-field Ca^2+^ imaging and local field potential recording in CA1. EC – extracellular electrode. (B) Resting GCaMP fluorescence and grid of ROIs used for analysis. (C) Time-aligned GCaMP fluorescence (ROIs indicated in B), LFP (band-pass-filtered at 0.5–100 Hz) and respiration/movement (resp) signal. Ca^2+^ clusters are indicated by gray, SPWs by red arrows. Note that CaTs are detected with or without a concurrent SPW. (D) Boxed regions marked in C at higher magnification. Note that SPW-related CaTs (+SPWs) display faster rise kinetics and higher amplitudes as compared to SPW-unrelated CaTs (–SPW). (E) Raster plots indicating CaT times per ROI from WT and KO^Emx1^ mice. Insets: Resting GCaMP fluorescence (F_0_) and binary area plots of Ca^2+^ clusters indicated on the left. (F) In NKCC1 KO^Emx1^ mice, mean CaT frequencies per ROI were significantly reduced. (G) Conditional NKCC1 deletion reduced the frequency of –SPW Ca^2+^ clusters. (H–J) +SPW and –SPW Ca^2+^ clusters display differences in rise kinetics (H), cluster area (I) and amplitude (J). SI – sampling interval. (K) Sample ΔF/F_0_ traces (mean of all analyzed ROIs) and time-aligned raster plots before and after local superfusion of gabazine (40 µM). Red arrows indicate SPWs. (L) Quantification of +SPW and –SPW Ca^2+^ cluster frequencies. (M) Sample raster plots demonstrating cluster activity in the visual neocortex as measured through the intact skull. Insets: Transcranial GCaMP fluorescence overlaid with binary area plots of three spatially confined cluster events indicated on the left (scale bars, 200 µm). (N–P) Quantification of the average CaT frequency per ROI (N), mean cluster area (O) and cluster frequency (P). Each open circle represents a single animal. Data are presented as mean ± SEM. ns – not significant, ^*^ P < 0.05, ^***^ P < 0.001. See also Figures S5 and S6 and Table S6.

For quantification, the field of view was subdivided into a regular grid of regions of interest (ROIs; Figure 6B). We found that the mean frequency of spontaneous CaTs per ROI was significantly lower in KO^Emx1^ mice than that in WT littermates (WT: 5.8 ± 0.3 min^−1^, n = 12, KO^Emx1^: 4.6 ± 0.4 min^−1^, n = 10, P = 0.015, two-sample t-test; Figure 6F and Table S6). Likewise, total Ca^2+^ cluster frequency was reduced from 7.9 ± 0.4 in WT to 6.3 ± 0.4 in KO^Emx1^ mice (main effect genotype: P = 8.8 × 10^−3^, two-way ANOVA; Table S6). Thus, our data identify an excitatory network effect of NKCC1 expressed by PCs in the neonatal CA1. While there was no significant interaction between the independent variables (genotype × SPW), we found a significant reduction in the frequency of –SPW (P = 0.010), but not in +SPW events (P = 0.72, two-sample t-tests with Bonferroni correction; Figure 6G). As compared to +SPW Ca^2+^ clusters, –SPW events had slower rise kinetics (Figure 6H), smaller cluster areas (Figure 6I) and lower CaT amplitudes (Figure 6J), indicating that they represent a separate class of CA1 network events. Interestingly, no genotype-dependent difference was found in any of these parameters (interaction genotype × SPW: at least P > 0.5, two-way ANOVAs; Figures 6H–6J and Table S6), demonstrating that network dynamics in the conditional absence of NKCC1 qualitatively resembled those observed in WT mice.

Ample theoretical and *in vitro* evidence indicates that depolarizing GABAergic transmission can mediate both excitation and inhibition in a context-dependent manner (Jean-Xavier et al., 2007; Kolbaev et al., 2011; Morita et al., 2006). We therefore examined the net contribution of endogenously released GABA by locally blocking GABA_A_Rs and found that superfusion of the hippocampal window with gabazine (40 µM) differentially influenced +SPW and –SPW Ca^2+^ clusters at P3–4. The frequency of +SPW Ca^2+^ clusters was massively augmented, while that of –SPW Ca^2+^ clusters was significantly decreased (Figures 6K and 6L). In addition, amplitudes of both event classes increased following GABA_A_R block, but this effect was significantly stronger for +SPW Ca^2+^ clusters (for statistical details, see #7 in Table S6).

Collectively, our data support the notion that GABAergic transmission facilitates the generation of –SPW Ca^2+^ clusters and additionally imposes inhibitory constraints, especially on +SPW events.

In contrast to our results obtained for the hippocampus, previous studies employing electrophysiological (Marguet et al., 2015; Minlebaev et al., 2007) or optical (Kirmse et al., 2015) measurements did not reveal a contribution of NKCC1 to spontaneous network activity in the neonatal neocortex. As this observation might imply brain region-dependent differences in the role of NKCC1 and, thus, depolarizing GABA (Murata et al., 2020), we next examined the consequences of a PC-specific NKCC1 disruption in the neonatal visual cortex (P3–4).

Cell-attached recordings in acute slices confirmed that puff application of a GABA_A_R agonist induced AP firing in PCs from WT, but not in PCs from KO^Emx1^, mice, demonstrating a loss of GABA-mediated excitation in the conditional absence of NKCC1 (Figures S6A–S6C). Moreover, total *Slc12a2* mRNA levels in the visual cortex of neonatal NKCC1 KO^Emx1^ mice were strongly reduced as compared to WT littermates (Figure S6D). Using *in vivo* wide-field Ca^2+^ imaging, we found that major aspects of spontaneous network activity were unaffected by NKCC1 loss, both at P3–4 (Figures 6M–6P) and P9–10 (Figures S6E–6H). Spindle bursts represent the main electrophysiological signature of Ca^2+^ clusters in the neonatal visual cortex (Kirmse et al., 2015). In agreement with data from optical measurements, multi-site LFP recordings from awake head-fixed P4 and P7 neonates confirmed that spindle burst properties including occurrence rates, lengths, and power spectral densities were unaltered in KO^Emx1^ mice (Figures S6I–6K).

Collectively, our data indicate that PC-specific expression of NKCC1 facilitates the generation of correlated network activity in a brain region- and event type-specific manner.

### Long-term alterations of in vivo hippocampal network dynamics upon NKCC1 deletion in Emx1-lineage cells

As correlated spontaneous activity is thought to be required for the proper maturation of neuronal circuits (Kirischuk et al., 2017; Leighton et al., 2016), we next investigated the long-term developmental effects of a PC-specific NKCC1 deletion onto *in vivo* hippocampal network dynamics in urethane-anesthetized 13-14 weeks-old mice. We recorded depth profiles of spontaneous LFPs along the CA1-dentate gyrus axis. Hippocampal LFPs from both WT and NKCC1 KO^Emx1^ mice showed paradoxical (REM) sleep-like activity, i.e., theta/gamma oscillations and non-REM slow wave sleep (SWS)-like LFP epochs with the characteristic SPW-ripple (SPW-R) complexes in CA1. These activity patterns alternated spontaneously throughout the recording session (Figure 7A). Quantitative LFP analysis of theta power depth profiles during REM-like epochs with similar theta phase depth profiles (see Methods and Figures S7A and S7B) revealed increased theta amplitudes in the hippocampal layers receiving direct input from entorhinal cortex [*stratum lacunosum moleculare* (s.l.m) and *stratum moleculare* (s.m.)], whereas amplitudes of low-gamma and multiunit activity frequency bands showed no significant changes in KO^Emx1^ mice (Figures 7B and 7C).

**Figure 7.**
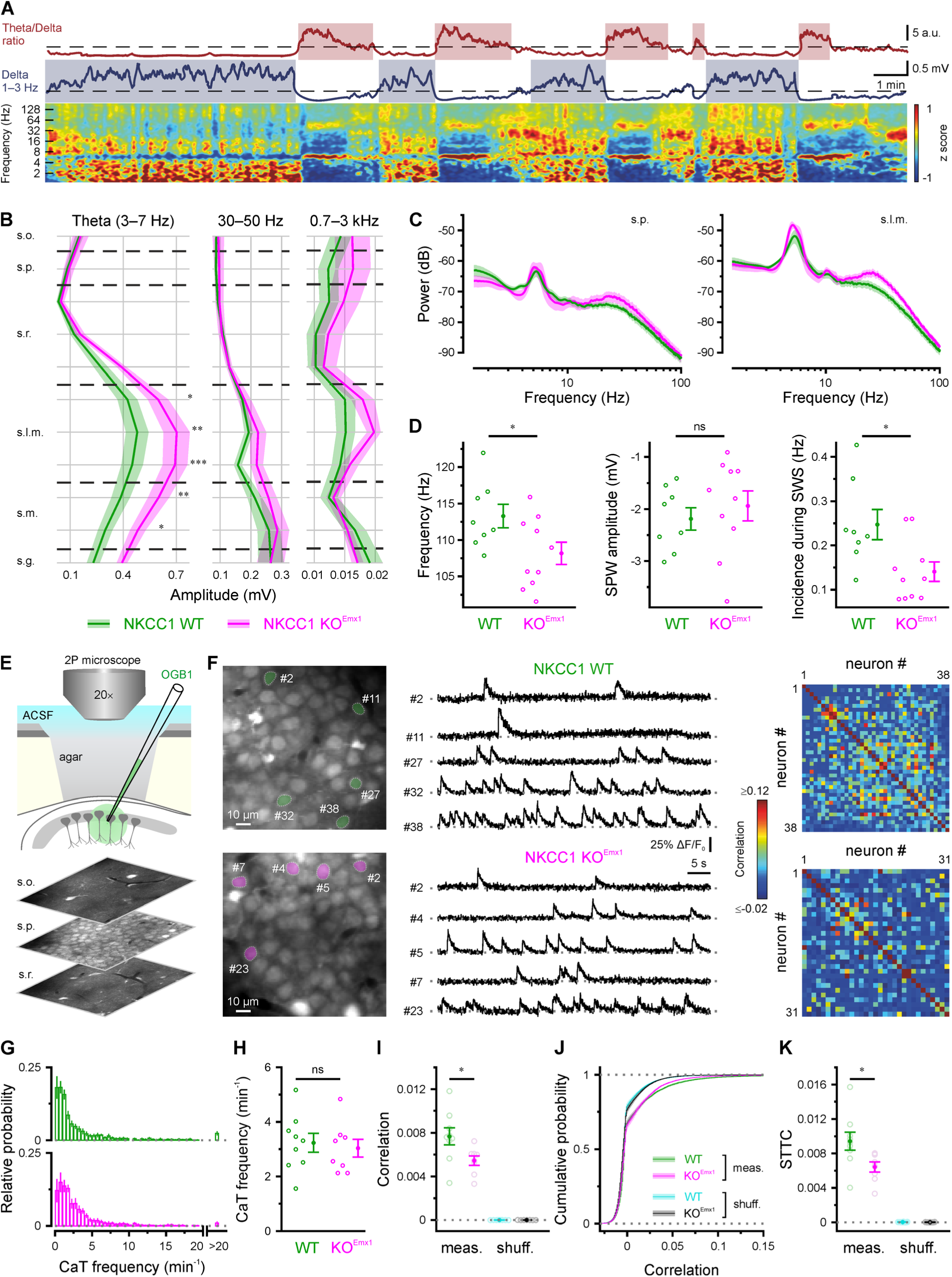
Long-term alterations of *in vivo* hippocampal network dynamics upon NKCC1 deletion in *Emx1*-lineage cells. (A) *Top*: Representative example of REM- and slow wave sleep- (SWS) like activity classification of the hippocampal LFP in adult urethane-anesthetized mice. SWS- and REM-like activity in s.l.m. showed characteristic inverted peaks in the delta band (blue trace) and theta/delta ratio (red trace), respectively. *Bottom*: Z-score normalized wavelet spectrogram showing the differences in theta and delta power between REM- and SWS-like oscillations. (B) Band-limited power analysis of the different strata in the dorsal hippocampus during REM-like activity. Theta amplitude (left) in s.l.m. and s.m. was higher in KO^Emx1^ compared to WT mice. Low-gamma (middle) and MUA (0.7–3 kHz, right) frequency bands were unaffected, as were the remaining frequency bands analyzed (see Methods). Shaded areas represent ± SEM across animals. (C) Power spectra comparison between s.p. and s.l.m. (D) SPW-R events occurring during SWS-like activity showed lower spectral frequency (left) and incidence (right) in KO^Emx1^ compared to WT mice. However, ripple-associated SPWs showed no significant amplitude difference between the genotypes (middle). (E) Experimental arrangement and sample OGB1 fluorescence images from s.o., s.p. and s.r. (F) *Left*: two-photon fluorescence images (OGB1); *middle*: sample ΔF/F_0_ traces; *right*: Pearson correlation matrices for all analyzed neurons obtained from fields of view shown on the left. (G) Distribution of CaT frequencies per cell (averaged across animals). (H) Mean CaT frequencies. (I) Mean pairwise Pearson correlation coefficients for measured data and randomly shuffled CaT trains. (J) Cumulative distributions of pairwise Pearson correlation coefficients. (K) Mean pairwise spike time tiling coefficients (STTC) for measured data and randomly shuffled CaT trains at identical mean frequencies. Each open circle represents a single animal. Data are presented as mean ± SEM. ns – not significant, ^*^ P < 0.05, ^**^ P < 0.01, ^***^ P < 0.001. See also Figure S7 and Table S7.

Analysis of SPW-Rs during SWS-like LFP epochs revealed lower ripple spectral frequencies, unchanged SPW amplitudes and decreased occurrence rates (Figure 7D). Cross-frequency coupling between theta and gamma oscillations, i.e., the modulation of gamma amplitudes by theta phase, was unaltered in KO^Emx1^ mice (Figures S7C and S7D). In addition, peak spectral frequencies in the theta (3–7 Hz) or gamma (30–100 Hz) range were similar in both genotypes (Figure S7E). Together, our data indicate that NKCC1 deletion from forebrain PCs altered the properties of theta oscillations in hippocampal layers receiving entorhinal input as well as intrahippocampal dynamics during ripple oscillations.

We further analyzed network dynamics at single-cell resolution using two-photon Ca^2+^ imaging in *stratum pyramidale* (s.p.) of mice (P45–50) under isoflurane anesthesia (Figure 7E). In line with published data (Busche et al., 2012), distributions of single-cell CaT frequencies were broad and strongly skewed to the right in either genotype (Figures 7F and 7G). Moreover, mean CaT frequencies per animal did not differ significantly between WT and KO^Emx1^ mice (Figure 7H). We further computed pairwise Pearson correlations based on binary CaT time courses convolved with a Gaussian kernel. We found that, by the age investigated, CA1 network activity had undergone a developmental transition to a largely desynchronized state in both genotypes (Rochefort et al., 2009). In quantitative terms, however, pairwise Pearson correlations were significantly lower in KO^Emx1^ mice as compared to their WT littermates (Figures 7I and 7J). Likewise, the spike-time tiling coefficient, a computationally independent affinity metric (Cutts et al., 2014), was significantly lower in the conditional absence of NKCC1 (Figure 7K). These genotype-dependent differences were abolished by randomly shuffling CaT times, confirming that they had not arisen from an inequality of CaT frequencies in individual cells.

Collectively, our data demonstrate subtle long-term alterations in both intrahippocampal dynamics and entorhinal cortex-hippocampus communication due to PC-specific NKCC1 deletion.

## Discussion

### Region- and event type-dependent contribution of NKCC1 to network dynamics in vivo

Recent studies revealed that NKCC1 mediates net chloride accumulation in immature neurons *in vivo*, giving rise to a mainly depolarizing action of GABA in the intact developing brain (Kirmse et al., 2015; Sulis Sato et al., 2017; van Rheede et al., 2015). However, the *in vivo* network functions of NKCC1-dependent GABAergic depolarization remained largely elusive (reviewed in Kirmse et al., 2018). In particular, systemic pharmacological approaches targeting NKCC1 yielded equivocal results (Marguet et al., 2015; Sipilä et al., 2006). By conditionally deleting *Slc12a2* from the vast majority of telencephalic glutamatergic cells, we demonstrate that NKCC1 facilitates synchronized network activity in the neonatal CA1 *in vivo*. Specifically, NKCC1 deletion led to a selective reduction in the frequency of –SPW Ca^2+^ clusters (by on average ∼20%), while leaving their kinetics, active area and amplitude unaffected. In contrast, SPWs and +SPW Ca^2+^ clusters were unaffected (Figures 5 and 6). Developmental compensations cannot be excluded from our data, but we emphasize that NKCC1 loss did reduce [Cl^−^]_i_ and profoundly attenuated network synchrony in acute slices. Additionally, no compensatory alterations in intrinsic or synaptic properties were detected that could explain the observed minor effect on network dynamics *in vivo*. Moreover, the impact of NKCC1 is region-specific, as its deletion failed to affect spindle bursts in the visual cortex, indicating that depolarizing GABAergic transmission is less important for driving network activity in neocortex. Conceptually, this conclusion is supported by a recent chemogenetic study, which revealed that the net action of GABA release is excitatory in CA1, but inhibitory in V1 (Murata et al., 2020). While the cellular and/or network mechanisms underlying the observed region specificity are currently unknown, it is worth noting that NKCC1 did contribute to the maintenance of excitatory GABA actions (i.e., [Cl^−^]_i_) in both CA1 (Figure 2) and the visual neocortex (Figure S6) *in vitro*.

Our data further revealed that +SPW and –SPW Ca^2+^ clusters were differentially modulated by local GABA_A_R inhibition. Specifically, gabazine substantially increased the frequency of +SPW Ca^2+^ clusters, but decreased that of –SPW events (Figure 6L). This opposite dependence on GABAergic transmission can possibly explain discrepant results of two well-designed recent studies, in which either inhibitory (Valeeva et al., 2016) or excitatory (Murata et al., 2020) effects of GABA were found to dominate. Hence, we argue that GABA_A_R signaling has a dual (excitatory-inhibitory) role in the neonatal CA1 circuitry. An interesting open question relates to the biophysical mechanism(s) underlying the excitatory *versus* inhibitory effects of GABA at this age. Ample theoretical and experimental evidence demonstrates that, even at the single-cell level, GABAergic depolarization may either increase or decrease neuronal firing in a manner that strongly depends on the activated GABA_A_R conductance in relation to E_GABA_ and the spatiotemporal patterning of synaptic inputs (Jean-Xavier et al., 2007; Kolbaev et al., 2011; Morita et al., 2006). Specifically, spindle bursts and SPWs are driven by highly synchronized glutamatergic inputs from the lateral geniculate nucleus (Ackman et al., 2012) and entorhinal cortex (Valeeva et al., 2019), respectively, while our LFP data suggest that this may be not the case with –SPW events.

Though speculative, such differences in synaptic inputs might explain why –SPW Ca^2+^ clusters are NKCC1-dependent, whereas spindle bursts and SPWs are not. It is hitherto unknown whether different classes of GABAergic interneurons (Pelkey et al., 2017) exert opposing network functions already in the neonatal hippocampus, as is the relative contribution of phasic *versus* tonic GABA_A_R signaling.

### Two classes of network events in the neonatal CA1 in vivo

Employing GCaMP-based mesoscopic Ca^2+^ imaging in combination with LFP recordings, we have identified the existence of two classes of network events in the neonatal CA1 *in vivo*. Due to space constraints, extracellular measurements were performed via a single electrode (placed in s.r.), which did not allow for reliable detection of HNOs as seen with multi-channel silicon probe recordings (Figure 5). Both +SPW and –SPW Ca^2+^ clusters were entirely AP-dependent (Figure S5), but differed in a number of basic parameters including rise kinetics, active area and amplitude (Figures 6H–6J). As compared to the neonatal visual cortex (Ackman et al., 2012; Kummer et al., 2016), Ca^2+^ clusters in CA1 generally exhibited a lower degree of spatial confinement. This was particularly prominent for +SPW events, which tended to activate virtually the entire field of view (Valeeva et al., 2020) and, in line with previous reports (Karlsson et al., 2006; Valeeva et al., 2019), frequently followed myoclonic limb/body twitches (Figure S5). Taken together, these observations suggest that SPWs may provide a pan-CA1 input signal representing, for example, somatosensory feedback from neonatal movements. Of note, –SPW events constituted about two thirds of all Ca^2+^ clusters detected in CA1 at P3–4 (Figure 6G). Since GCaMP signals reflect neuronal firing rather than sub-threshold membrane fluctuations (Kerr et al., 2005), our data indicate that a considerable fraction of APs in *Emx1*^+^ CA1 PCs is not driven by cortically triggered SPWs (Valeeva et al., 2019). This conclusion is supported by previous electrophysiological data from freely moving rat pups (P4–6) demonstrating that ∼60% of CA1 MUA bursts were not associated with SPWs (Leinekugel et al., 2002). Some –SPW Ca^2+^ clusters followed +SPW events with short delays and, hence, might reflect *tails* of MUA described previously (Leinekugel et al., 2002; Marguet et al., 2015). However, based on descriptive differences (see above) and their differential dependence on NKCC1 (Figures 5 and 6G) and GABA_A_R signaling (Figure 6L), we argue that –SPW events are mechanistically distinct. Based on their differential NKCC1 dependence, we propose that GDPs do not represent the *in vitro* analog of SPWs recorded *in vivo*. Since (I) – SPW events resemble GDPs in their pharmacological profile and (II) GDPs are generated even in the deafferented hippocampus *in vitro* (Menendez de la Prida et al., 1998), it is tempting to speculate that –SPW Ca^2+^ clusters might emerge from and reflect a property of local intrahippocampal circuits, whereas SPW generation requires cortical input (Valeeva et al., 2019).

### Developmental functions of NKCC1-dependent GABAergic depolarization

We provide evidence that the maturation of a number of passive and active (Figures 4A–4E) membrane properties were unaffected by the chronic absence of NKCC1 in PCs. This suggests that compensatory changes in intrinsic excitability, previously observed in a conventional NKCC1 knockout line (Sipilä et al., 2009) or after E15–P7 pharmacological NKCC1 blockade (Wang et al., 2011), may be due to loss of NKCC1 function in other cell types such as GABAergic interneurons, or reflect secondary alterations related to NKCC1 loss outside the brain. Furthermore, several basic characteristics of AMPAR- and GABA_A_R-mediated synaptic transmission were unaltered in NKCC1 KO^Emx1^ mice (Figures 4F–4I). This observation challenges the hypothesis of the importance of NKCC1-dependent GABAergic depolarization as a universal requirement for the NMDA receptor (NMDAR)-dependent developmental unsilencing of glutamatergic synapses (reviewed in Kirmse et al., 2018; Wang et al., 2008, 2011) – despite the fact that depolarizing GABA may facilitate NMDAR-mediated currents by attenuating their voltage-dependent Mg^2+^ block (Leinekugel et al., 1997). Collectively, our data indicate that network effects induced by NKCC1 disruption in telencephalic PCs directly result from changes in E_GABA_ (i.e., [Cl^−^]_i_). Thus, we also confirm a large body of *in vitro* (Dzhala et al., 2005; Sipilä et al., 2006; Yamada et al., 2004) and recent *in vivo* (Kirmse et al., 2015; Sulis Sato et al., 2017; van Rheede et al., 2015) evidence indicating that NKCC1 maintains a relatively high [Cl^−^]_i_ in immature neurons. Of note, NKCC1 deletion did not completely abolish depolarizing responses to GABA in acute slices, suggesting that factors other than NKCC1 contribute to chloride accumulation in developing hippocampal PCs (Pfeffer et al., 2009). In addition, optical and electrophysiological measurements in adult mice demonstrated relatively subtle alterations of network dynamics in NKCC1 KO^Emx1^ mice (Figure 7) that were not associated with overt performance deficits in several hippocampus-dependent behavioral tasks (Figure 1). These findings may also be of clinical relevance, as NKCC1 has emerged as a potential target for the treatment of neurodevelopmental disorders (Deidda et al., 2015; Marguet et al., 2015; Tyzio et al., 2014).

While our study does not rule out more specific long-term developmental functions of NKCC1 and/or roles of NKCC1 under pathophysiological conditions, our data converge to suggest that basal physiological NKCC1 activity is largely dispensable for several key aspects of cortical network maturation.

## Acknowledgments

We thank Ina Ingrisch and Sindy Beck for excellent technical assistance and Dr. John Dempster (University of Strathclyde, Glasgow) for adapting the software WinFluor. This work was supported by the Priority Program 1665 (HO 2156/3–1/2 to K.H., KI 1816/1–1/2 to K.K., KI 1638/3–1/2 to S.J.K., IS63/5–1 to D.I. and HU 800/8–1/2 to C.A.H.), the Collaborative Research Center/Transregio 166 (B3 to K.H., K.K.), the Collaborative Research Center 1089 (A05 to D.I., S.L.M.) and the Research Unit 3004 (KI 1816/5-1) of the German Research Foundation, the Federal Ministry of Education and Research (NEURON ACRoBAT to C.A.H., 01GQ0923 to K.H. and O.W.W.), and the Interdisciplinary Centre for Clinical Research Jena (K.K., K.H.).

## Author Contributions

K.K., K.H., C.A.H., S.J.K. and D.I. designed research with contributions from all authors. J.G., C.Z., S.L.M, T.H., T.F., R.H., M.G. and K.K. performed experiments. K.K., J.G., C.Z., S.L.M., T.H., T.F., V.R., M.G., C.F., A.U., R.M.N. and D.I. analyzed data. All authors contributed to the interpretation of the data. K.K., J.G., C.Z., K.H., S.L.M., D.I., C.A.H. and S.J.K. wrote the manuscript with contributions from all authors.

## Methods

### Animals

A detailed list of mouse strains used in this study is provided in Table S8. All animal procedures were performed with the approval of the local governments (Thüringer Landesamt für Verbraucherschutz, Bad Langensalza, Germany, and Landesamt für Natur-, Umwelt- und Verbraucherschutz, NRW, Recklinghausen, Germany) and complied with European Union norms (Directive 2010/63/EU). Animals were housed in standard cages with 14h/10h (Jena) or 12h/12h (Cologne) light/dark cycles. *Emx1^IREScre^* (stock no. 005628), *GCaMP3^LSL^* (Ai38, stock no. 014538), *tdTomato^LSL^* (Ai14, stock no. 007908) and C57BL/6J (stock no. 000664) mice were originally obtained from the Jackson Laboratory. *NKCC1^flox^* (*Slc12a2^flox^*) mice were generated as previously described (Antoine et al., 2013). The age of the animals is indicated in the Results section. Double transgenic animals were obtained by crossing *Emx1^IREScre/wt^:NKCC1^flox/flox^* female to *NKCC1^flox/flox^* male mice. Triple transgenic animals were obtained by crossing *NKCC1^flox/wt^:GCaMP3^LSL/LSL^* or *NKCC1^flox/wt^:tdTomato^LSL/LSL^* male to *Emx1^IREScre/IREScre^:NKCC1^flox/wt^* female mice. Animals having the genotype *Emx1^wt/wt^:NKCC1^flox/flox^, Emx1^IREScre/wt^:NKCC1^wt/wt^:GCaMP3^LSL/wt^* or *Emx1^IREScre/wt^:NKCC1^wt/wt^:tdTomato^LSL/wt^* are collectively referred to as WT, and animals with *the genotype Emx1^IREScre/wt^:NKCC1^flox/flox^, Emx1^IREScre/wt^:NKCC1^flox/flox^:GCaMP3^LSL/wt^* or *Emx1^IREScre/wt^:NKCC1^flox/flox^:tdTomato^LSL/wt^* are collectively referred to as KO^Emx1^. Mice of either sex were used. Experiments were performed blinded to genotype.

### Preparation of acute brain slices

Animals were decapitated under deep isoflurane anesthesia. The brain was quickly removed and transferred into ice-cold saline containing (in mM): 125 NaCl, 4 KCl, 10 glucose, 1.25 NaH_2_PO_4_, 25 NaHCO_3_, 0.5 CaCl_2_, and 2.5 or 6.0 MgCl_2_, gassed with 5% CO_2_ /95% O_2_ (pH 7.4). Horizontal or coronal brain slices (350 µm) were cut on a vibratome and stored for at least 1 h before their use at room temperature in artificial cerebrospinal fluid (ACSF) containing (in mM): 125 NaCl, 4 KCl, 10 glucose, 1.25 NaH_2_PO_4_, 25 NaHCO_3_, 2 CaCl_2_, and 1 MgCl_2_, gassed with 5% CO_2_ /95% O_2_ (pH 7.4). For recordings, slices were placed into a submerged-type recording chamber on the microscope stage (Nikon Eclipse FN1, Nikon Instruments Inc.), which was equipped with near-infrared differential interference contrast video-microscopy (ACSF flow rate ∼3–4 ml min^−1^). All experiments were performed at ∼32 °C.

### Patch-clamp recordings in vitro

Electrophysiological signals were acquired using an Axopatch 200B or Multiclamp 700B amplifier, a 16-bit AD/DA board (Digidata 1440A or Digidata 1550A) and the software pClamp 10 (Molecular Devices). Signals were low-pass filtered at 1–3 kHz and sampled at 10–20 kHz. To estimate E_GABA_ non-invasively while minimizing the risk of unintended membrane breakthrough, GABA-induced membrane potential (V_m_) alterations were measured using low-concentration gramicidin perforated-patch current-clamp recordings (I = 0 mode of the amplifier) (Perkins, 2006; Zhang et al., 2019; Zhu et al., 2008). Recording pipettes (5–8 MΩ) were filled with the following solution (in mM): 140 K-gluconate, 1 CaCl_2_, 2 MgCl_2_, 11 EGTA, 10 HEPES (pH 7.25), supplemented with 40–80 µg/ml gramicidin. Measured voltages were corrected for the calculated liquid junction potential (∼16 mV). Recordings were performed in the presence of antagonists of ionotropic glutamate receptors (10 µM DNQX, 50 µM APV) and voltage-gated Na^+^ (0.5 µM TTX) and Ca^2+^ (100 µM CdCl_2_) channels so as to block amplification of voltage changes by intrinsic conductances. A saturating puff of the GABA_A_R agonist isoguvacine (100 µM, 2 s) was applied twice to the soma of the recorded cell at an interval of 2 min. As demonstrated before, V_peak_ approximates E_GABA_ under these conditions (Zhu et al., 2008).

For measurements of spontaneous or agonist-induced action currents, tight-seal cell-attached recordings were performed in voltage-clamp mode using glass pipettes filled with the following solution (in mM): 150 NaCl, 4 KCl and 10 HEPES (pH 7.4). In addition, 50 µM Alexa Fluor 555 was added to the pipette solution to detect potential membrane breakthrough. Holding current was manually zeroed before each experiment.

For recording sGPSCs, the intra-pipette solution contained (in mM): 150 Cs-methanesulfonate, 5 NaCl, 10 HEPES, 5 EGTA, 0.5 CaCl_2_ (pH 7.3). Holding potential was set to 0 mV (compensated for the calculated liquid junction potential of ∼10 mV) to isolate sGPSCs from glutamatergic sEPSCs. sGPSCs were detected using template matching (pClamp 10).

GABA_A_R-mediated mGPSCs and AMPA receptor-mediated mEPSCs were recorded using the whole-cell patch-clamp technique. Intra-pipette solution contained (mM) (1) for mGPSCs: 145 CsCl, 5 NaCl, 10 HEPES, 2 QX-314, 0.2 EGTA, 2 Mg-ATP, 0.3 Na-GTP (pH 7.3), or (2) for mEPSCs: 125 Cs-methanesulfonate, 20 CsCl, 5 NaCl, 10 HEPES, 2 QX-314, 0.2 EGTA, 2 Mg-ATP, 0.3 Na-GTP (pH 7.3). Alexa Fluor 555 (50 µM) was regularly included for morphological identification of the recorded cell. Pipette resistance was 3–5 MΩ when filled with the solution mentioned above. Holding potential was set to −70 mV (not corrected for liquid junction potentials). Access resistance was monitored by applying hyperpolarizing pulses of 10 mV. Only recordings with an access resistance below 30 MΩ were accepted. Series resistance compensation was not applied.

To measure passive properties and intrinsic excitability, whole-cell voltage-clamp and current-clamp recordings were used, respectively. Intra-pipette solution contained (in mM): 133 K-gluconate, 12 KCl, 5 NaCl, 10 HEPES 10, 0.2 EGTA, 2 Mg-ATP, 0.3 Na-GTP (pH 7.3), regularly supplemented with Alexa Fluor 488 (10 µM). Ionotropic glutamate and GABA receptor antagonists (10 µM DNQX, 50 µM APV, 10 µM bicuculline methiodide [BMI]) were added to the ACSF to abolish recurrent excitation and to minimize synaptic noise. In voltage-clamp recordings, holding potential was set to –70 mV; in current-clamp measurements, membrane potential was biased to –70 mV before a series of 500-ms episodic current injections (from –120 pA to 120 pA with 10 pA increments) was applied. Only recordings with an access resistance below 30 MΩ were accepted. Series resistance compensation was not applied.

### Confocal Ca^2+^ imaging in vitro

Large-scale Ca^2+^ imaging was performed in horizontal cortico-hippocampal slices from mice conditionally expressing the genetically encoded Ca^2+^ indicator GCaMP3 in *Emx1*-positive cells. For single-cell Ca^2+^ imaging, cells were loaded with the membrane-permeable Ca^2+^ indicator Oregon Green 488 BAPTA-1 AM (OGB1) using multi-cell bolus-loading in s.p. of hippocampal CA1 or CA3, as indicated in the Results section. Fluorescence signals were acquired at ∼22.5 Hz using a CSU10 Nipkow-disc scanning unit (Yokogawa Electric Corp.) in combination with a Rolera XR FAST 1394 CCD camera (12 bit, QImaging Corp.) and the software Winfluor 3.7.5 (Dr. John Dempster, University of Strathclyde, Glasgow) or Streampix 5 (NorPix). Excitation light at 488 nm was provided by a single wavelength solid-state laser (Sapphire CDRH-LP, Coherent) via a 2.5×/0.075 (Zeiss) or 16×/0.8 W (Nikon) objective for large-scale and single-cell imaging, respectively. An acousto-optic tunable filter (GH18A, Gooch & Housego) was used to adjust the final laser power of excitation.

### Surgical preparation, anesthesia and animal monitoring for in vivo imaging

Animals were placed onto a warm platform and deeply anesthetized with isoflurane (3.5% for induction, 1–2% for maintenance) in pure oxygen (flow rate: 1 l/min). The skin overlying the skull was disinfected and locally infiltrated with 2% lidocaine (s.c.). Eyes of adult mice were lubricated with a drop of eye ointment (Vitamycin) and covered with aluminum foil. Scalp and periosteum were removed, and a custom-made plastic chamber with a central borehole (Ø 2.5–4 mm) was fixed on the skull using cyanoacrylate glue (for CA1 recordings at P3–4: 3.5 mm rostral from lambda and 1.5 mm lateral from midline; at P45–50: 2.5 mm caudal from bregma and 2.2 mm lateral from midline; for V1 recordings: 1.5–2.5 mm lateral from midline, immediately rostral to the transverse sinus).

For the hippocampal window preparation (Mizrahi et al., 2004), the plastic chamber was tightly connected to a preparation stage and subsequently perfused with ACSF containing (in mM): 125 NaCl, 4 KCl, 25 NaHCO_3_, 1.25 NaH_2_PO_4_, 2 CaCl_2_, 1 MgCl_2_ and 10 glucose (pH 7.4, 35–36°C). A circular hole was drilled into the skull using a tissue punch (outer diameter 1.8 mm for P3–4 and 2.3 mm for P45–50 mice). The underlying cortical tissue and parts of corpus callosum were carefully removed by aspiration using a vacuum supply and a blunt 27G or 30G needle. Care was taken not to damage alveus fibers. As soon as bleeding stopped, the animal was transferred to the microscope stage. For transcortical LFP recordings in CA1 (see Figure S5), a small craniotomy was performed above the somatosensory cortex (4 mm rostral from lambda and 1.5 mm lateral from midline) using a 30G needle.

For visual cortex measurements, the mouse was transferred to the microscope stage, and the plastic chamber perfused with ACSF (as above). At P9–10, a craniotomy was performed above the left occipital cortex using a 27G needle. Care was taken not to damage the underlying dura mater. At P3–4, measurements were performed through the intact skull.

During *in vivo* recordings, body temperature was continuously monitored and maintained at close to physiological values (34–37°C) by means of a heating pad and a temperature sensor placed below the animal. Spontaneous respiration was monitored using a differential pressure amplifier (Spirometer Pod and PowerLab 4/35, ADInstruments). For wide-field Ca^2+^ imaging at P3–4 and P9–10, isoflurane was discontinued after completion of the surgical preparation and gradually substituted with the analgesic-sedative nitrous oxide (up to the fixed final N_2_O/O_2_ ratio of 3:1, flow rate: 1 l/min). Experiments started 60–120 min after withdrawal of isoflurane. For two-photon Ca^2+^ imaging experiments at P45–50, the isoflurane concentration was gradually reduced to 0.4–0.6%. At the end of each experiment, the animal was decapitated under deep isoflurane anesthesia.

### In vivo wide-field Ca^2+^ imaging and LFP recording

The recording chamber was continuously perfused with ACSF (as above). In case of recordings in CA1, a tungsten microelectrode (Tunglass-1, 0.8 MΩ impedance, Kation Scientific) was lowered just above the hippocampal formation. ACSF was then removed and the hippocampal window was filled up with agar (1%, in 0.9 mM NaCl). As soon as the agar solidified, the chamber was reperfused with ACSF. Next, the microelectrode was slowly lowered to 200–250 µm below the hippocampal surface preparation. The final electrode depth was determined when the recorded SPWs showed a polarity reversal.

For transcortical LFP recordings in CA1 (Figure S5), a tungsten microelectrode was lowered to 1400–1600 µm below the skull. The final electrode depth was determined when recorded SPWs qualitatively showed a polarity reversal. In order to confirm the proper electrode position *post hoc*, an electrolytic lesion was performed via the electrode (4 V for 5 s, Master-8, A.M.P.I.) at the end of the experiment. Electrophysiological signals were acquired and low-pass-filtered at 3 kHz using an EXT-02F/2 amplifier (npi electronic), a 16-bit AD/DA board (PowerLab 4/35, ADInstruments) and the software LabChart 8 (ADInstruments). Signals were and sampled at 20 kHz.

Wide-field epifluorescence Ca^2+^ imaging was performed in mice conditionally expressing GCaMP3 in *Emx1*-positive cells. One-photon excitation was provided by either a xenon arc lamp (Lambda LS, Sutter Instrument) coupled via a liquid light guide to the epifluorescence port of a Movable Objective Microscope (Sutter Instrument) for V1 recordings or a 470-nm collimated LED (Thorlabs) connected to a Zeiss Axioskop for hippocampal recordings and filtered at 472/30 nm (AHF Analysentechnik). Emission was separated from excitation light at 495 nm and long-pass-filtered at 496 nm (AHF Analysentechnik). Images were acquired using a 10×/0.3 NA water immersion objective lens (Zeiss) and a 12-bit Rolera-XR camera (QImaging) operated by the software Streampix 5 (NorPix) or Winfluor 3.7.5 (Dr. John Dempster, University of Strathclyde, Glasgow). Using 4×4 hardware binning (174×130 pixels), the sampling rate was ∼74.3 Hz. For V1 recordings the field of view was 1,134×867 µm and for hippocampal recordings 877×696 µm. Recording time per condition typically amounted to 20–30 min.

### In vivo two-photon Ca^2+^ imaging in adult mice

The recording chamber was continuously perfused with ACSF (as above). Cells were loaded with the membrane-permeable Ca^2+^ indicator Oregon Green 488 BAPTA-1 AM (OGB1) using multi-cell bolus-loading in s.p. of hippocampal CA1 (Stosiek et al., 2003). ACSF was then removed and the hippocampal window was filled up with agar (1%, in 0.9 mM NaCl) and covered with a cover glass. As soon as the agar solidified, the chamber was reperfused with ACSF. To achieve de-esterification, recordings started ∼60 min after OGB1 injection. Imaging was performed using a Movable Objective Microscope (Sutter Instrument) equipped with two galvanometric scan mirrors (6210H, MicroMax 673XX Dual Axis Servo Driver, Cambridge Technology) and a piezo focusing unit (P-725.4CD PIFOC, E-665.CR amplifier, Physik Instrumente) controlled by a custom-made software written in LabVIEW 2010 (National Instruments) (Kummer et al., 2015) and MPScope (Nguyen et al. 2006). Fluorescence excitation at 800 nm was provided by a tuneable Ti:Sapphire laser (Chameleon Ultra II, Coherent) using a 20×/1.0 NA water immersion objective (XLUMPLFLN 20XW, Olympus). Emission light was separated from excitation light using a 670-nm dichroic mirror (670 DCXXR, Chroma Technology), short-pass filtered at 680 nm and detected by a photomultiplier tube (12 bit, H10770PA-40, Hamamatsu). Data were acquired using two synchronized data acquisition devices (NI 6110, NI 6711, National Instruments). Sampling rate was set 31.4 Hz (110×110 pixels, 106×106 µm). For each animal, spontaneous activity was recorded within 3–5 fields of view, each one usually for ∼20 min.

### Acute in vivo silicon probe depth recordings of awake head-fixed neonatal mice

*In vivo* depth recordings with linear 32-site silicon probes were performed in awake, head-fixed animals. For analgesia, animals received buprenorphine (0.025 mg/kg body weight subcutaneously [s.c.]) 30 min prior to anesthesia induction, which was done with 4% isoflurane in 100% oxygen in an induction chamber. Anesthesia was maintained using 1.0– 2.0% isoflurane throughout the entire surgical procedure. Once sufficient depth of anesthesia was achieved, which was verified by observing the absence of tail pinch and pedal withdrawal reflexes, a midline skin incision was performed on the top of the skull. For additional analgesia, skin incisions were treated with the local anesthetic bupivacaine (Bucain®-Actavis 0.25%). The periosteum was denatured by short (3–4 s) H_2_O_2_ (10%) treatment and subsequently removed with a cotton swab. The skull surface was rinsed with NaCl 0.9% solution and dried. For a common ground and reference electrode, a hole was drilled above the cerebellum. A silver wire was inserted above the dura of the cerebellum and fixed with dental cement. Subsequently, a metal tube was positioned over the cerebellar skull and attached with dental cement to facilitate fixation of the animal using the ear bars of the mouse stereotaxic apparatus (Stoelting). Mice were resting during the entire recording on a heated, custom-made padded brass block allowing unrestrained limb movements and, thereby, reducing movement artifacts in the recorded LFP. Constant ambient nest temperature of approximately 34°C was maintained on top of the brass block by use of a homeothermic heating pad (Stoelting), which was placed below the block and connected with a feedback sensor to the temperature controller. To insert silicon probes targeting the dorsal hippocampal CA1 or the primary visual (V1) cortex regions, burr holes were placed at the following stereotaxic coordinates: For dorsal CA1, rostro-caudal half of the lambda-bregma distance; dorso-ventral 1.9–2.3 mm, 1.2 mm lateral from the sagittal suture at P5, 1.4 mm at P7, and 1.65 mm at P12. For V1 recordings of all ages, the electrode positions were 0.1–0.2 mm anterior of lambda, 2.4-2.8 mm lateral and 0.8 mm dorso-ventral. Isoflurane anesthesia was discontinued once surgery was complete. Linear 32-site (A1x32-5mm-50-703-A32; NeuroNexus) or 16-site (A1x16-3mm-50-703-A16; NeuroNexus) silicon probes with a distance of 50 µm between the recording sites and a site surface of 703 µm², were implanted perpendicular to the brain surface along the hippocampal CA1-DG axis or in V1, respectively. The silicon probes were connected to a 1x preamplifier (Neuralynx) mounted to the stereotaxic instrument (Stoelting). Data were digitally filtered (0.5–9000 Hz bandpass) and digitized as 16-bit integers with a sampling rate of 32 kHz using a Digital Lynx 4S data acquisition system (Neuralynx). Neuralynx files were pre-processed into open formats and downsampled for neonatal LFP analysis to 1280 Hz using NDManager Plugins from Neurosuite (neurosuite.sourceforge.net). For unit analyses, files were high-pass filtered with a median filter (window size 39 samples), and common median referenced by subtracting the median of six neighboring channels (excluding nearest neighbors) in Matlab. Spikes were extracted for channels surrounding the CA1 s.p. by NDManager Plugins and the first three PCA components clustered by KlustaKwik2 (https://github.com/kwikteam/klustakwik2). Artefacts and low-quality units were removed by hand using Klusters (Neurosuite) and the resulting units merged into MUA for further analysis. Animal movements were detected by a piezoelectric sensor placed under the animal’s chest and recorded in parallel with the same sampling rate. Data acquisition was started about 15–30 min after probe insertion when recording conditions were stable. Probe positions were marked by the silicon probe tips labeled with the fluorescent dye DiI (Life Technologies) and verified in DAPI-stained (DAPI Fluoromount-GTM fluorescent; Southern Biotech) coronal slices, which, in combination with the depth profile of the LFP, allowed the post-hoc layer identification.

### In vivo hippocampal recordings in adult mice

Adult mice were initially anesthetized with 1.0–1.3 mg/g bodyweight urethane (Sigma, 10% w/v, in NaCl 0.9%). As it takes for urethane about 30 min to guarantee sufficient anesthesia after injection, mice were additionally anaesthetized with the inhalative anesthetic isoflurane for the first 30 min of the procedure (1.0–1.5% in 100% oxygen). Animals were placed in a stereotaxic apparatus where they remained head-fixed during the entire recording (Stoelting). Body temperature was kept stable at 36.5°C during the operation and the recording using a homeothermic heating pad (Stoelting). A midline skin incision was made on the top of the skull. For a common ground and reference electrode, a hole was drilled above the cerebellum and a stainless steel screw, connected to the ground wire, was inserted (lambda -1.5 mm, 1.0 mm lateral). For the hippocampal silicon probe, a 0.8-mm wide burr hole was placed at 2.0 mm posterior to bregma and 1.6 mm left to the midline. The surgical procedure lasted 25–30 min, and isoflurane anesthesia was stopped once the surgery was finished. A linear 32-site silicon probe with a distance of 50 µm between the recording sites (A1x32-5mm-50-177; NeuroNexus Technologies) was vertically inserted into the dorsal hippocampus (2.1 mm depth) along the CA1-DG-axis. Data acquisition, storage, and pre-processing for offline analysis were done as above.

### Behavioral experiments

#### Open field

Ten-week-old mice were placed in the middle of a white plastic box (50×50×50 cm), and spontaneous activity was recorded for 10 min using a CCD camera. The periphery was defined as the region outside of a 25×25 cm central square. The total distance covered and the time spent in each area were analyzed with TSE VideoMot2 software (TSE Systems).

#### Morris water maze

Spatial learning and memory were assessed in a circular pool (diameter 120 cm, height 60 cm) filled with opaque water (23 ± 1°C) to a depth of approximately 30 cm. In order to escape from the water, mice had to locate a hidden platform (diameter 10 cm) placed approximately

0.5 cm below the water surface. A fixed array of extra-maze cues (e.g. geometric shapes, cupboards) was available for spatial navigation. On the day before training, mice were given a single 60-s trial in the pool without a platform to habituate them to the test situation. During the next eight days, mice were tested for four trials each day in the hidden-platform task: Mice were individually released into the water at one out of four cardinal points, facing the pool wall. The release points changed pseudo-randomly across trials and days but were the same for all animals. If a mouse failed to locate the platform within 60 s, it was gently guided to it. Mice were allowed to rest on the platform for 20 s and returned to a warm cage for an inter-trial-interval of 30 s. 24 hours after the last training trial, a 60-s probe trial was implemented to assess spatial reference memory. To this end, the platform was removed from the pool, and mice started from the quadrant opposite to the former platform position. 24 hours after the second probe trial, a cued test (four 60-s trials) was carried out to control for non-mnemonic factors (e.g. visual or motor disturbances, motivation) that may influence task performance. Here, the platform was visible (raised 0.5 cm above the water surface and marked with a flag) and moved to a new position each trial. All trials were recorded by an automated video tracking system (TSE VideoMot2, TSE) to determine the path length (distance) during training and cued trials, and the percent time spent in each quadrant in the probe trials.

#### Fear conditioning

The apparatus (UgoBasile) consisted of four sound-attenuating cubicles, each containing an inner chamber that could be made distinct between trials by changing its shape, wall patterns, floor, illumination and ventilation. Mouse behaviour was monitored by a USB video camera connected to a PC running ANY-maze software (Stoelting). During acquisition, mice were placed into a transparent cuboid conditioning chamber with a shock grid floor that had 5 lx illumination, fan speed set at 100% and was cleaned with 70% ethanol. After 3 min, a tone was presented (conditioned stimulus (CS); 20 s, 80 dB, 9000 Hz) that co-terminated with an electric foot shock (unconditioned stimulus (US); 2 s, 0.7 mA). 60 s later, animals were returned to their home cages. To evaluate auditory-cued fear memory, mice were placed into a novel context (plexiglass cylinder, walls covered by a vertical pattern of black and white stripes, solid white plastic floor, cleaned with 5 % acetic acid, fan speed 50 %) and the house light was switched on (2 lx). Three minutes later, mice were re-exposed to the tone for 3 min, followed by another 60 s before they were returned to their home cage. Contextual fear was tested on the same day by re-exposing mice to the original conditioning context for 3 min. Freezing behaviour was analyzed offline from the video recordings.

#### Y maze

The apparatus consisted of three white, opaque plastic arms (40×6.5×10 cm) interconnected at an angle of 120°. A separate cohort of mice was used. Mice were placed into the center of the maze and allowed to freely explore until they completed 27 transitions. Entering an arm with all four paws was considered as transition. Movements were recorded by an automated video tracking system (TSE VideoMot2, TSE). The sequence of arm entries was analyzed, and triads (consecutive entries into three different arms) were counted to calculate the percentage of spontaneous alternations, defined as the ratio of the actual-to-possible completed triads: alternations (%) = [(number of completed triads)/(total number of arm entries – 2)] x 100.

#### Neonatal reflexes

Sensorimotor reflex development was assessed using the righting reflex and cliff avoidance tests. Mice were tested at postnatal days 4, 7 and 12, and scored twice with an intertrial interval of 30 sec and maximum trial duration of 30 sec. The righting reflex was assessed by placing mice on their back and measuring the time needed for righting themselves. Righting was considered successful when mice touched the ground with all four paws. Cliff avoidance reaction was measured by placing mice with their nose above the edge of the table. The time needed to turn the body 90° relative to the edge of the table was recorded.

#### Immunohistochemistry

*Emx1^IREScre/wt^:NKCC1^wt/wt^:tdTomato^LSL/wt^* (n = 3) and *Emx1^IREScre/wt^:NKCC1^flox/flox^:tdTomato^LSL/wt^*(n = 3) mice (P54–63) were deeply anaesthetized with isoflurane and transcardially perfused with 4% paraformaldehyde (PFA) in phosphate-buffered saline (PBS). Brains were post-fixed in 4% PFA overnight and sequentially cryoprotected in PBS containing 10% and 30% sucrose, respectively. Coronal sections (40 μm) were prepared on a cyromicrotome (HM400, Microm). For immunohistochemistry, slices were washed in PBS, permeabilized with 0.2% Triton X-100 and blocked with 3% normal donkey serum (NDS). For primary antibodies raised in mice, unspecific binding was minimized by pre-incubation with Fab fragments (donkey anti-mouse) for 2h. Free-floating sections were incubated overnight with primary antibodies in PBS containing 3% NDS/0.2% Triton X-100 at 4 °C. Following rinsing, secondary antibodies were applied for 3 h at room temperature in PBS supplemented with 3% NDS/0.2% Triton X-100. Slices were rinsed again and mounted on slides using Fluoromount (SouthernBiotech). A detailed list of antibodies used in this study is provided in Table S8. DAPI was applied to counterstain nuclei. Fluorescence images (z-stacks with 1-μm vertical separation of optical sections) were acquired with an LSM 710 laser-scanning confocal microscope and a 40× oil immersion objective (Zeiss).

### Quantitative PCR

Mice (P2–3) were decapitated under deep isoflurane anesthesia. Brains were quickly removed and transferred to an ice-cold preparation platform under a stereomicroscope. From each animal, either the hippocampi or the left visual cortex were manually dissected, frozen and stored at –80°C. Total RNA was isolated by the phenol chloroform extraction method and reverse transcribed into cDNA by the use of the RevertAid First Strand cDNA Synthesis Kit (ThermoFisher). The qPCR was performed as previously described (Frahm et al., 2017) using specific primers for *Slc12a2* (forward [fw] primer: TCAGCCATACCCAAAGGAAC; reverse [rv] primer: AACACACGAACCCACAGACA). For normalization, *Gapdh* (fw primer: CAACAGCAACTCCCACTCTTC; rv primer: GGTCCAGGGTTTCTTACTCCTT) and *Hmbs* (fw primer: GTTGGAATCACTGCCCGTAA; rv primer: GGATGTTCTTGGCTCCTTTG) were used as reference genes.

### Chemicals

A detailed list of chemicals used in this study is provided in Table S8.

### Analysis and quantification

#### Electrophysiological recordings in vitro

To quantify the irregularity of sGPSCs, a (frequency-dependent) metric referred to as burst index was defined as the fraction of inter-sGPSC intervals ≤50 ms. Only cells with ≥50 detected sGPSCs were included in the analysis. Burst indices computed for measured data were compared to burst indices reflecting a Poisson point process in which sGPSCs occur randomly. In such a case, inter-event intervals are exponentially distributed, and the probability of an event within a time t is given by:

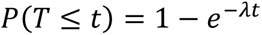

where *λ* is the mean event rate (in Hz) and t = 0.05 s.

In current-clamp recordings, action-potential threshold was determined as the V_m_ value at which the first time derivative of V_m_ exceeded 10 V/s (Stimfit). mGPSCs and mEPSCs were analyzed using Minianalysis 6.0.

#### Confocal Ca^2+^ imaging in vitro

To correct for x-y drifts during measurements, image stacks were registered using a template-matching algorithm (ImageJ plugin by Q. Tseng). Fluorescence signals were obtained from manually selected regions of interest and expressed as relative changes from resting fluorescence levels (ΔF/F_0_). For large-scale imaging, onsets and offsets of CaTs were manually determined using a threshold criterion, considering changes in ΔF/F_0_ as an event if they exceeded two times the standard deviation of the baseline for at least 200 ms. The area under the curve of CaTs was computed for all regions and slices in which at least five events were detected. For single-cell imaging, Ca^2+^ transients (CaTs) were detected using a template-matching algorithm implemented in pClamp 10 (Molecular Devices), and their times of peak were defined as the times of CaT occurrence. Network events were operationally defined as GDPs as follows: (1) CaTs were classified as GDP-related if they fell into a 500-ms-long time interval during which the fraction of active cells was ≥20%. (2) Neighboring GDP-related CaTs were assigned to the same GDP if both shared a 500-ms-long time interval during which the fraction of active cells was ≥20%, otherwise to separate GDPs. For each GDP, active cells were ranked according to their CaT times, and GDP half-width was calculated as the difference in CaT times corresponding to the 25^th^ and 75^th^ percentiles of active cells. Slices with less than 10 GDPs were excluded from analysis.

#### In vivo widefield Ca^2+^ imaging and LFP recording

For analysis, periods of movement artefacts were identified by visual inspection of raw image sequences and discarded because they frequently resulted in obviously false-positive detection results. Image sequences were then binned (10×10), resulting in a regular grid of 17×13 = 221 ROIs (ROI dimensions: for V1: 66.7×66.7 µm, for hippocampus: 51.6×51.6 µm). For CA1 data, we excluded those ROIs from further analysis that overlapped with the extracellular microelectrode or minor blood residues. Next, light intensity-versus-time plots were extracted using ImageJ. Unbiased peak detection of spontaneous CaTs was performed using a template-matching algorithm implemented in pClamp 10. ‘Clusters’ of CaTs were defined on a temporal basis. (I) For CA1: If two neighboring CaTs were separated by less than 300 ms, they were considered to belong to the same cluster, otherwise to two separate clusters. Only clusters with a fraction of active ROIs ≥10% of all analyzed ROIs were considered. (II) For the visual cortex: If two neighboring CaTs were separated by less ≤10 frames (∼135 ms), they were considered to belong to the same cluster, otherwise to two separate clusters. The mean frequencies of CaTs per ROI and the mean cluster frequencies were calculated as the ratio of number of events per corrected recording time (i.e., total recording time minus cumulative movement artefact-associated recording time). Amplitudes of CaTs were calculated as relative changes from resting fluorescence levels, defined for each individual event. Maximum rise slopes were extracted from the first derivative of raw fluorescence traces following Savitzky–Golay smoothing (2^nd^ order, 8 sampling intervals), and normalized to F_0_. Amplitudes and rise slopes were averaged for each Ca^2+^ cluster. The cluster area was calculated as the ratio of the number of active ROIs *versus* the total numbers of ROIs analyzed. For hippocampal recordings, SPWs were detected as follows: Raw data (20 kHz) were first down-sampled by a factor of four and filtered at 0.5–100 Hz (pClamp 10). Candidate events were detected using a threshold algorithm. Visual inspection of high-pass filtered (>100 Hz) signals was used to exclude false-positive events. Detected SPWs were separated into two either (I) movement-associated or (II) non-movement-associated SPWs based on the respiration/movement signal from a pressure sensor placed below the mouse. A Ca^2+^ cluster was assigned to a SPW if the first cluster-related CaT was within the interval [SPW time – 100 ms, SPW time + 400 ms].

#### In vivo two-photon Ca^2+^ fluorescence imaging

Somatic fluorescence signals were obtained from manually selected regions of interest and expressed as relative changes from resting fluorescence levels (ΔF/F_0_). CaTs were detected using UFARSA, an acronym for *Ultra-Fast Accurate Reconstruction of Spiking Activity* (Rahmati et al., 2018). Reconstructed CaT times were translated into a binary vector for each trace. Pairwise Pearson correlation coefficients were determined for all possible cell pairs analyzed in a given image sequence using custom routines (Matlab, MathWorks). To this end, binary CaT vectors were first convolved with a Gaussian kernel with a standard deviation of two sampling intervals. Spike-time tiling coefficients (STTCs) were computed for all possible cell pairs with a synchronicity window of three sampling intervals (∼96 ms) using custom routines (Matlab, MathWorks) (Cutts et al., 2014). Pearson correlation coefficients and STTCs derived from measured data were compared to those from simulated data obtained by randomly shuffling (uniform distribution; 4,000 times) CaT times of all cells. When performing this randomization, the mean CaT frequency of each cell was kept unchanged.

#### Acute in vivo hippocampal & V1 depth recordings of unanesthetized neonatal mice

Analyses were performed using custom-written functions and scripts (Matlab, MathWorks). The CA1 s.p. was identified as the channel with minimum coherence to nearest neighbors in the frequency range of 8–64 Hz and confirmed to be the LFP reversal, corrected if necessary, using Neuroscope (Neurosuite). Current source density was computed as the second spatial derivate of the 1–200 Hz forward and reverse 2-pole Butterworth-filtered LFP. Hippocampal HNOs and V1 spindle bursts were marked using 10–30 Hz (HNOs) or 5–25 Hz (V1 spindle bursts) power increases in the multitaper spectrogram (Chronux) of the whitened LFP by three blinded individual observers who corrected each other’s results. SPWs in neonates lack ripples, and were detected on the median-filtered 10 Hz high-passed CSD and LFP using a novel algorithm. Briefly, peaks in the inverted CSD with half-width between 4–50 ms exceeding the 99^th^ percentile CSD negativity in prominence, and no closer than 12 ms apart, are detected using Matlab findpeaks() for each channel. Solitary peaks without a neighboring (±2 channels) CSD peak in a ±14 ms window, or smaller than -0.1 mV in a ±3 ms window of the LFP, are then discarded. Finally, the largest peak in each 7 ms window is kept, and any putative SPW following another by less than 40 ms (typically, the rebound sink in s.o. for strong radiatum / lacunosum SPWs) is merged with the previous SPW. SPWs with the most negative CSD sink above and including the reversal layer are considered ‘oriens’ SPWs, those below as ‘rad/lm’ (radiatum / lacunosum). For statistics, amplitude was taken from the most negative LFP channel in a window ± 3 ms around the CSD-derived SPW trough. HNO spectrograms displayed were computed using Morse wavelets with a time-bandwidth product of 30, normalized to median absolute deviation (MAD) units per frequency bin, and then smoothed with a 15 ms wide Hamming window. MUA was smoothed with a 7 ms Hanning window for display.

#### Acute in vivo dorsal hippocampal depth recordings of urethane-anesthetized mice

All analyses of hippocampal LFP were made using custom-written Matlab scripts. REM- and SWS-like activity in recordings of urethane-anesthetized mice were classified by thresholding the delta band (1–3 Hz) and theta (3–7 Hz)/delta ratio. The threshold was manually set for each recording. Delta and theta bands were computed as the absolute value of the Hilbert transform of the bandpass filtered LFP and smoothed with a 10-sec quadratic kernel. Although we used a low dose of urethane (1.0–1.3 mg/g) in these experiments, we observed a high variability in several electrophysiological markers, such as theta phase shifts, theta power and cross frequency coupling, between the REM-like periods within a recording session. Urethane is known to disrupt theta rhythm generation in the entorhinal cortex by blocking local NMDA receptors (Gu et al., 2017). In order to reduce this variability, we calculated the theta phase shift (Figure S7A) between s.p. and s.l.m for each REM-like period and concentrated subsequent analyses on those periods showing shift values within the highest quartile across a recording session. The resulting phase shift values were closer to those observed in awake condition (Figure S7B) (Lubenov et al., 2009). Band-limited power (BLP) analysis of the different strata in the dorsal hippocampus was performed for the following frequency bands: delta, 1–3 Hz; theta, 3–7 Hz; sigma, 7–16 Hz; beta, 16–30 Hz, low-gamma, 30–50 Hz; mid-gamma, 40–70 Hz; high-gamma, 70–100 Hz; high frequency oscillation (HFO) 1, 110–170 Hz; HFO2, 140–240 Hz; HFO3, 300–600 Hz and MUA, 700–3000 Hz. First, the original LFP at 32 kHz was downsampled to 1280 Hz (except for the MUA band). Then this downsampled signal was bandpass filtered, rectified and resampled to 250 Hz. Hippocampal strata identification was assessed based on stereotaxic and electrophysiological markers such as the laminar distribution of the aforementioned BLPs during REM- and SWS-like activity, theta-gamma coupling as quantified by the modulation index (MI) (Tort et al., 2008), ripple amplitude, theta phase shifts (Lubenov et al., 2009) and spiking activity. All recordings were post-hoc histologically verified. Ripples were detected by bandpass filtering the CA1 s.p. recording site at 90–180 Hz and thresholding at 4 SD of the absolute value of the Hilbert transform. Following detection, the frequency and duration of individual ripple events were calculated from time-resolved generalized Morse wavelets spectrograms of ±30 ms peri-threshold crossing windows. For the MI laminar distribution and comodulogram analyses, we selected the recording site with highest theta amplitude (s.l.m.) as reference to extract the theta phases. The tested amplitude-modulated frequency bands were low-, mid-, high-gamma and HFO.

### Statistical analysis

Statistical analyses were performed using OriginPro 2018, SPSS Statistics 22/24, Statistica 13.3, and Microsoft Excel 2010. Unless otherwise stated, the statistical parameter n refers to the number of (I) animals for *in vivo* experiments, (II) slices for *in vitro* Ca^2+^ imaging and (III) cells in patch-clamp recordings, respectively. All data are reported as mean ± standard error of the mean (SEM), if not stated otherwise. The Kolmogorov–Smirnov test or Shapiro–Wilk test was used to test for normality of the data. Parametric testing procedures were applied for normally distributed data; otherwise non-parametric tests were used. In the case of two-sample t-tests and unequal group variances, Welch’s correction was applied. For multi-group comparisons, analysis of variance (ANOVA) was applied. Analysis of covariance (ANCOVA) was used to control for covariate effects. P values (two-tailed tests) < 0.05 were considered statistically significant. Details of the statistical tests applied are provided in Tables S1–S7.

### Data and code availability

All datasets generated during this study are available from the corresponding author upon request.

