## Supplemental Information for "Intraneuronal chloride accumulation via NKCC1 is not essential for hippocampal network development *in vivo*"

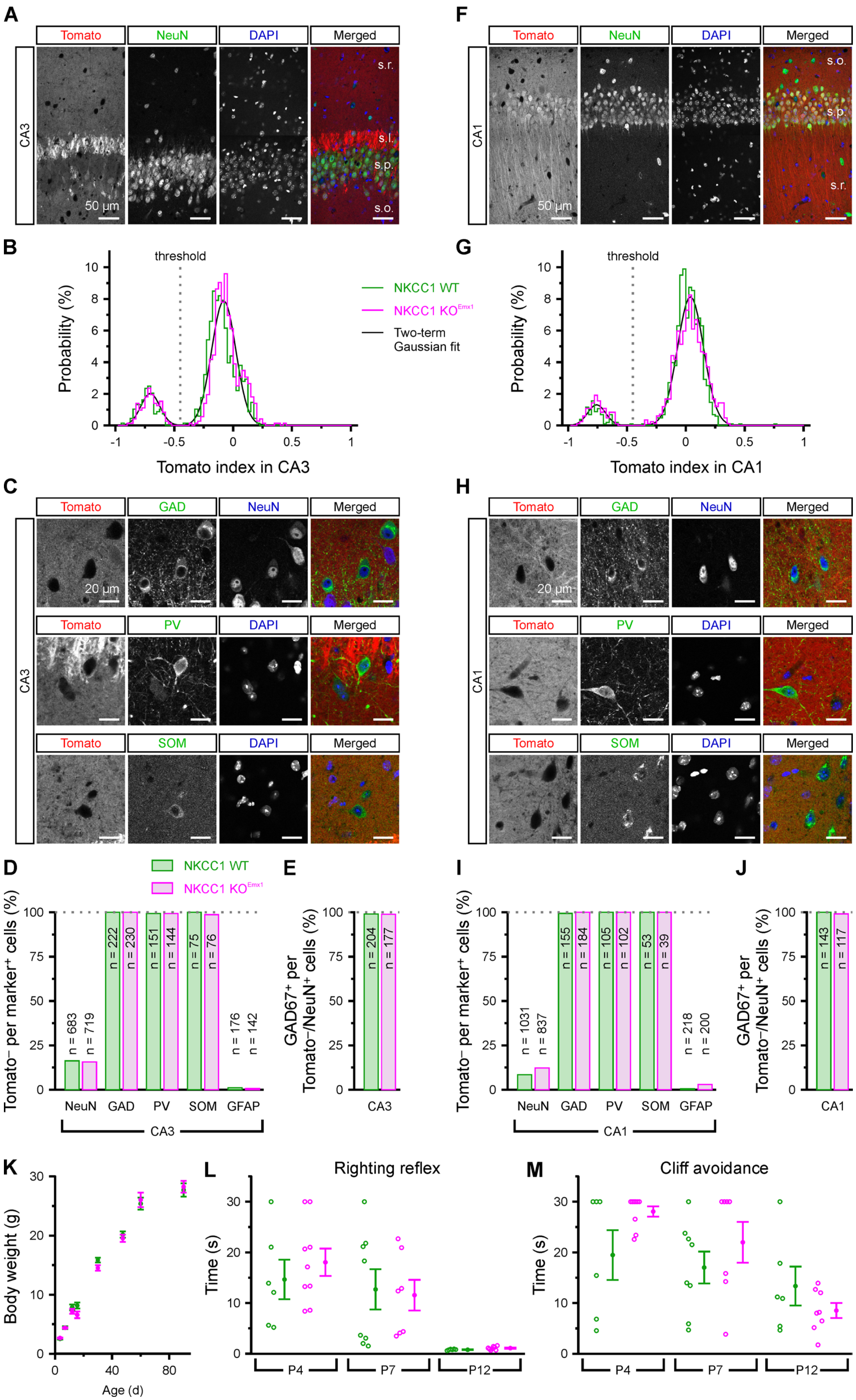

A
Tomato
NeuN
DAPI
Merged
CA3
s.r.
s.l.
s.p.
s.o.
50 µm
F
Tomato
NeuN
DAPI
Merged
CA1
s.o.
s.p.
s.r.
50 µm
B
threshold
Probability (%)
Tomato index in CA3
NKCC1 WT
NKCC1 KO^Emx1
Two-term Gaussian fit
G
threshold
Probability (%)
Tomato index in CA1
C
Tomato
GAD
NeuN
Merged
20 µm
Tomato
PV
DAPI
Merged
Tomato
SOM
DAPI
Merged
CA3
H
CA1
D
NKCC1 WT
NKCC1 KO^Emx1
Tomato− per marker+ cells (%)
n = 683
n = 719
n = 222
n = 230
n = 151
n = 144
n = 75
n = 76
n = 176
n = 142
NeuN
GAD
PV
SOM
GFAP
CA3
E
GAD67+ per Tomato−/NeuN+ cells (%)
n = 204
n = 177
CA3
I
Tomato− per marker+ cells (%)
n = 1031
n = 837
n = 155
n = 184
n = 105
n = 102
n = 53
n = 39
n = 218
n = 200
NeuN
GAD
PV
SOM
GFAP
CA1
J
GAD67+ per Tomato−/NeuN+ cells (%)
n = 143
n = 117
CA1
K
Body weight (g)
Age (d)
L
Righting reflex
Time (s)
P4
P7
P12
M
Cliff avoidance
Time (s)
P4
P7
P12

**Figure S1. Related to Figure 1.** Fate mapping of *Emx1*-lineage cells in hippocampal areas CA3 and CA1. (A) Confocal images of a histological section demonstrating *Emx1*$^{IREScre}$-driven tdTomato expression in area CA3 of an adult *Emx1*$^{IREScre}$*:NKCC1*$^{wt/wt}$*:tdTomato*$^{LSL}$ (WT) mouse. Note that only a minor fraction of neurons (NeuN$^{+}$) lacks tdTomato. Also note strong tdTomato signal of mossy fibers in *stratum lucidum* (s.l.). (B) tdTomato$^{+}$ and tdTomato$^{-}$ cells can be reliably differentiated on the basis of a fluorescence index (see Methods) in both WT and *Emx1*$^{IREScre}$*:NKCC1*$^{flox/flox}$*:tdTomato*$^{LSL}$ (KO$^{Emx1}$) mice. (C) Confocal images of histological sections demonstrating that glutamate decarboxylase 67 (GAD), parvalbumin (PV) or somatostatin (SOM) are exclusively present in tdTomato$^{-}$ cells. (D) Quantification of tdTomato$^{-}$ cells among marker$^{+}$ cells. Note that virtually all GABAergic (GAD67$^{+}$) cells lack tdTomato. (E) Quantification of GAD67$^{+}$ cells among tdTomato$^{-}$ neurons (NeuN$^{+}$). Note that virtually all tdTomato$^{-}$ neurons express GAD67. Three mice per genotype were included in the analysis. Numbers refer to the numbers of cells analyzed. (F–J) Analogous to panels A–E, but based on data acquired in area CA1. (K) Body weight during postnatal development. (L) Righting reflex: Time needed by the pup to flip onto its feet from a supine position. (M) Cliff avoidance: Time needed by the pup to turn, i.e. move paws and snout away from the edge. A synopsis of the applied statistical procedures can be found in Table S1.

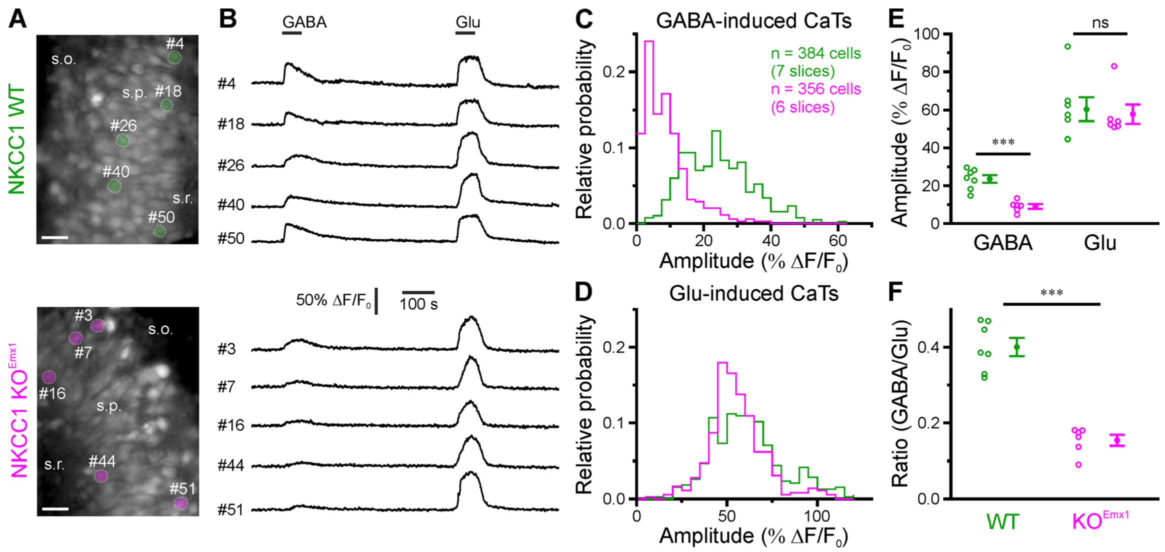


**Figure S2. Related to Figure 2.** NKCC1 deletion in *Emx1*-lineage cells attenuates GABA-induced somatic $Ca^{2+}$ transients. (A) Fluorescence images of OGB1-stained CA3 cells in acute hippocampal slices. Scale bars, 20 µm. (B) Sample $Ca^{2+}$ traces from cells highlighted in A. Experiments were performed in the continuous presence of TTX (0.5 µM). GABA (100 µM) and glutamate (Glu, 100 µM) were bath applied for 60 s. (C–D) Distributions of amplitudes of GABA- and Glu-induced CaTs. (E) Mean amplitudes of GABA- and Glu-induced CaTs per slice. (F) Mean ratios of amplitudes of GABA- vs. Glu-induced CaTs. Each open circle represents a single slice. Data are presented as mean ± SEM. ns – not significant, *** $P < 0.001$. A synopsis of the applied statistical procedures can be found in Table S2.

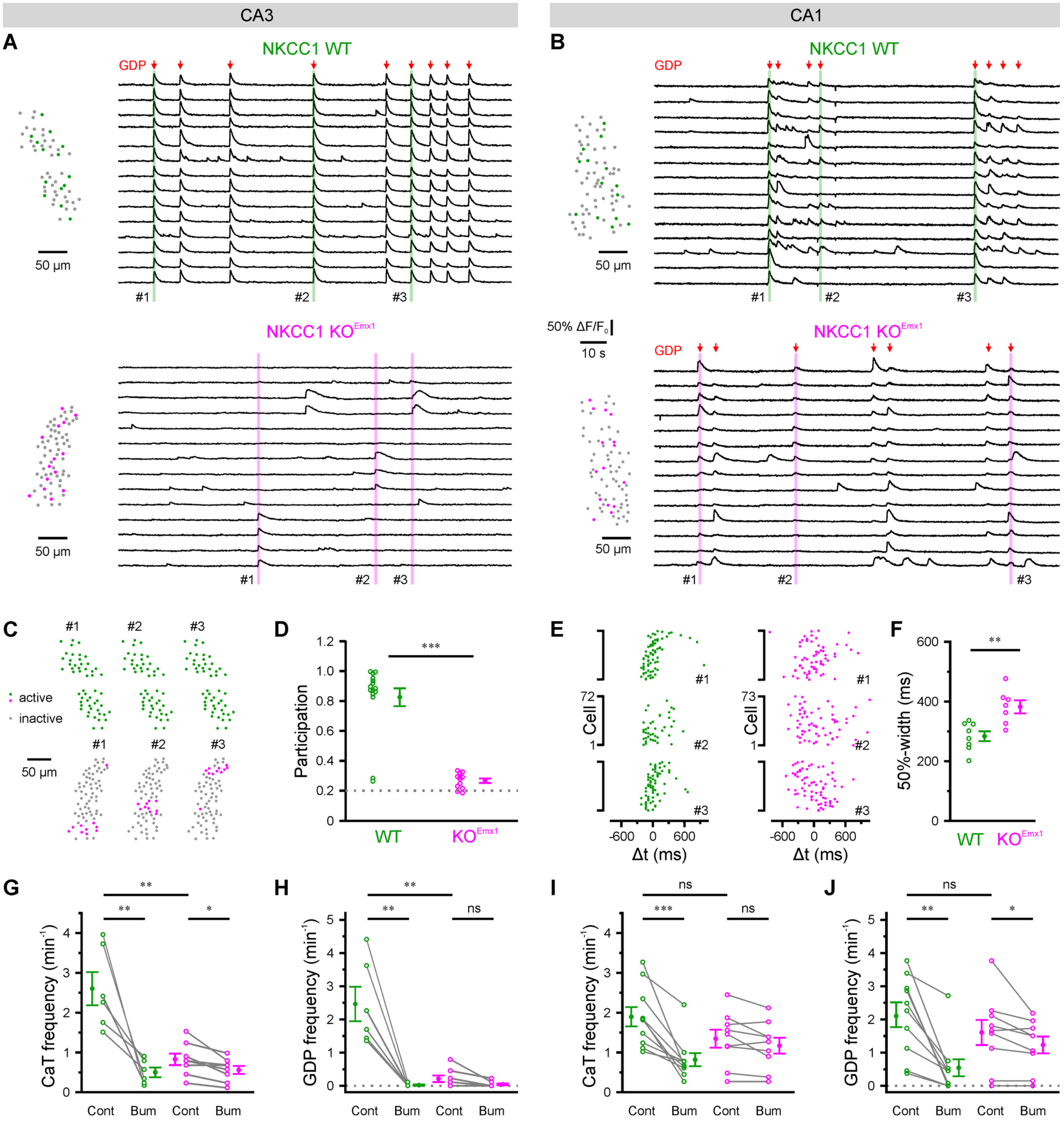


**Figure S3. Related to Figure 3.** Single-cell $Ca^{2+}$ imaging reveals differential effects of conditional NKCC1 deletion on network dynamics in CA3 and CA1 *in vitro*. (A–B) Sample $Ca^{2+}$ traces of OGB1-stained neurons in s.p. of CA3 (A) or CA1 (B). Left: Relative position of all analyzed neurons. Colored dots indicate the neurons selected for sample $Ca^{2+}$ traces. Red arrows indicate network events with ≥ 20% active neurons (GDPs; for definition see Methods). (C) Activity maps of network events highlighted in A. Note that network events in KO$^{Emx1}$ CA3 are restricted to a small fraction of analyzed neurons. (D) Fraction of active neurons within GDPs, averaged for all GDPs per slice. (E) Temporal distribution of CaTs around the median time point of example GDPs highlighted in B. Note that in KO$^{Emx1}$ CA1, GDPs show a higher temporal jitter. (F) Mean 50%-widths of GDPs. (G–J) Frequencies of CaTs and GDPs in the absence (Cont) or presence (Bum) of the NKCC1 antagonist bumetanide (50 µM) in CA3 (G–H) and CA1 (I–J). (D, F–J) Each open circle represents a single slice. Data are presented as mean ± SEM. ns – not significant, $^{*}$ $P < 0.05$, $^{**}$ $P < 0.01$ $^{***}$ $P < 0.001$. A synopsis of the applied statistical procedures can be found in Table S3.

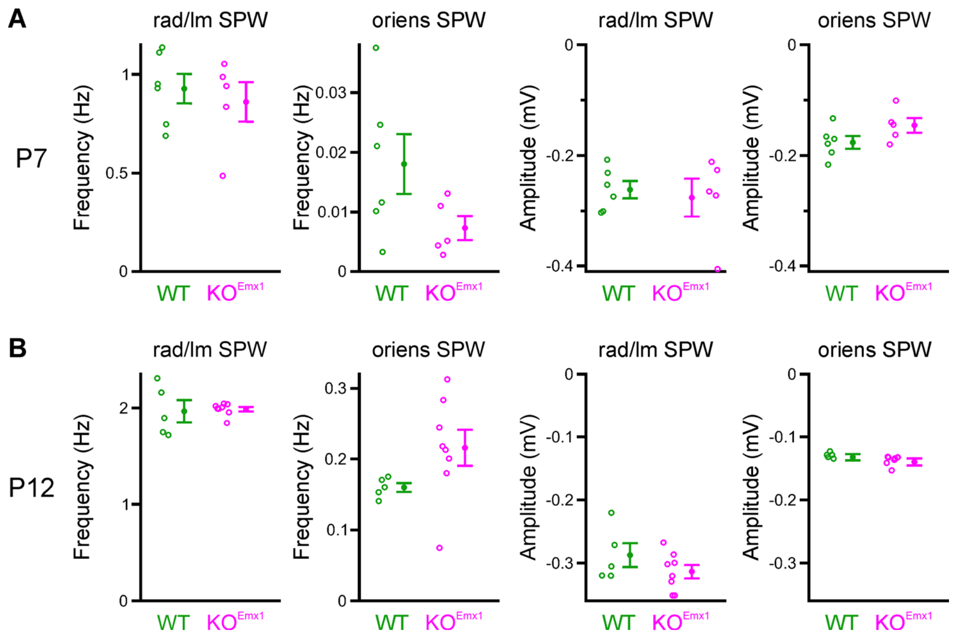


**Figure S4. Related to Figure 5.** Hippocampal sharp waves *in vivo* at P7 and P12 are unaltered by the conditional absence of NKCC1. (A) Neither SPWs with sinks below s.p. ("rad/lm SPW") or above ("oriens SPW") differ in occurrence frequency or amplitude (minimum trough negativity in the LFP) at P7. (B) As in (A), the $KO^{Emx1}$ animals show similar SPW parameters to WT at P12. A synopsis of the applied statistical procedures can be found in Table S5.

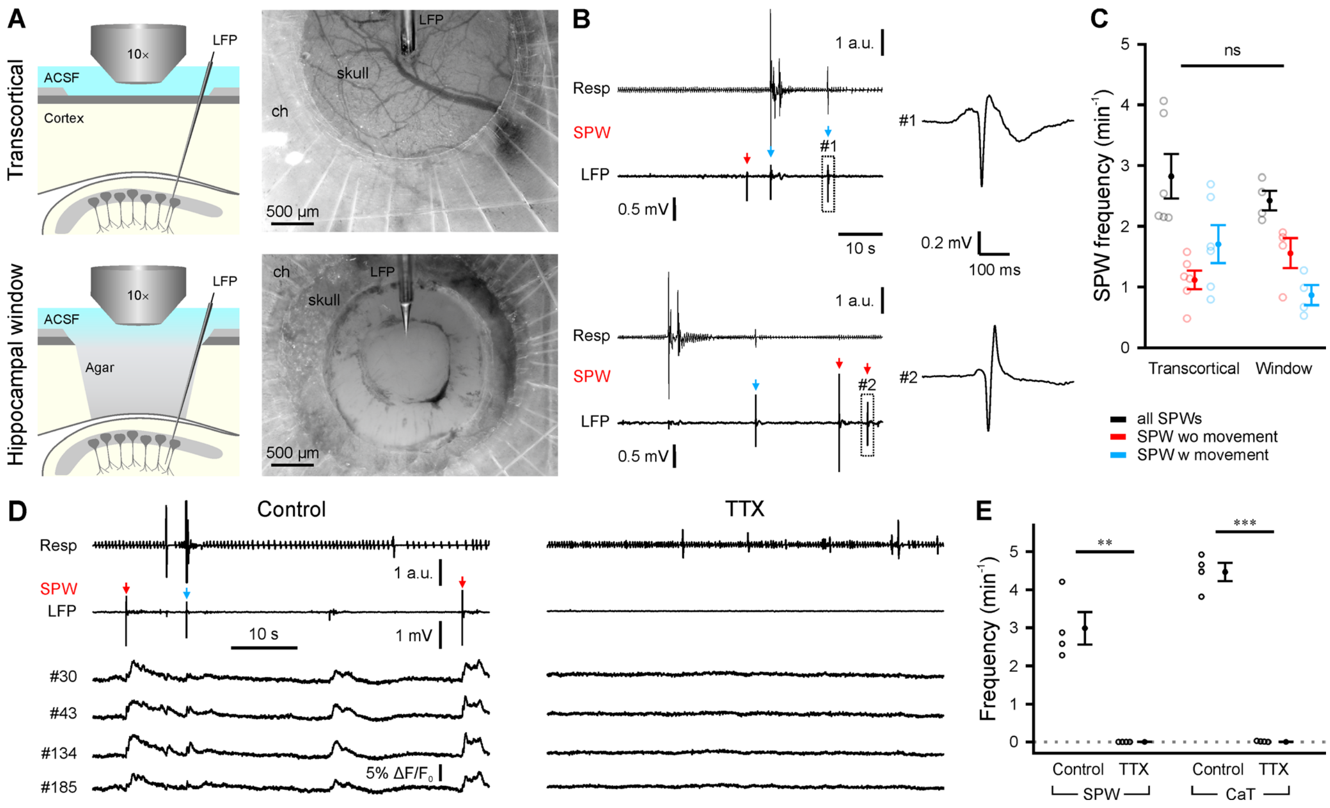


**Figure S5. Related to Figure 6.** Sharp waves (SPWs) are unaffected by focal cortex aspiration and depend on action-potential firing. (A) Experimental arrangements and brightfield images of transcortical and hippocampal window recordings. ch – recording chamber. (B) *Left*: Sample LFP recording (0.5–100 Hz) and respiration/movement signal (Resp) for both recording conditions. Note that SPWs (arrows) were either associated (blue) or not associated (red) to brief animal movements. *Right*: Two SPWs are shown at higher temporal resolution. (C) Mean frequency of all SPWs (black), movement-associated SPWs (blue) and not movement-associated SPWs (red). Data were obtained from C57BL/6J mice (unpaired). (D) Sample LFP recording and $Ca^{2+}$ traces time-aligned to respiration/movement signal under control conditions and after local superfusion of TTX (5 µM). Note that both CaTs and SPWs were blocked by TTX. (E) Mean frequencies of SPWs and CaTs. Data were obtained from one WT and three $KO^{Emx1}$ animals. Each open symbol represents a single animal. Data are presented as mean ± SEM. ns – not significant, $^{**}$ $P < 0.01$. A synopsis of the applied statistical procedures can be found in Table S6.

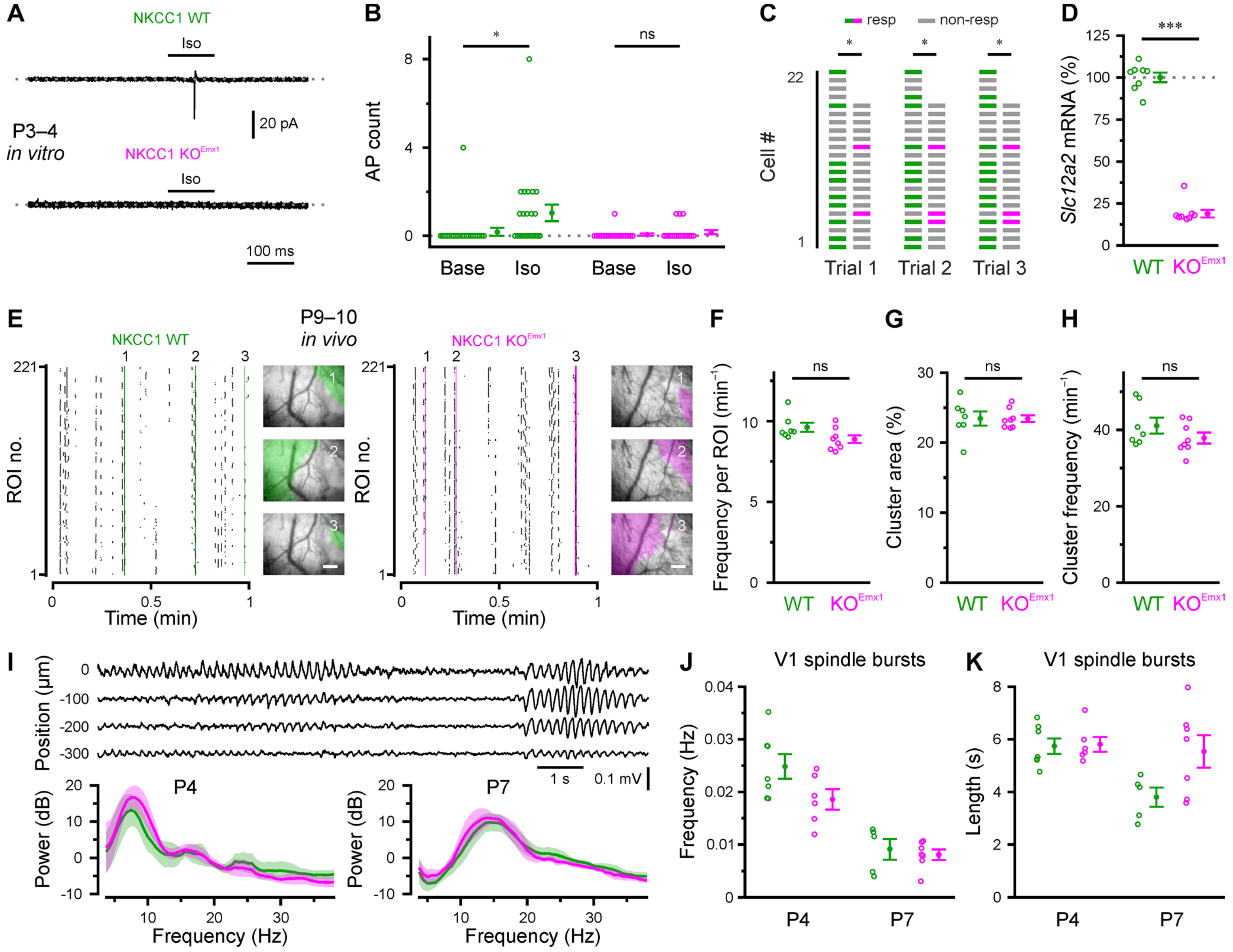


**Figure S6. Related to Figure 6.** NKCC1 deletion in *Emx1*-lineage cells does not affect network activity in the visual cortex before eye-opening. (A) Sample cell-attached recordings in response to puff-application of isoguvacine. (B) Number of action currents detected in 300-ms intervals immediately before (Base) and after (Iso) puff onset. Each open symbol represents a single cell. (C) The fraction of responsive cells (resp) was significantly lower for $KO^{Emx1}$ as compared to WT mice. (D) Total visual cortex *Slc12a2* mRNA levels normalized to the geometric mean of WT. (E) Sample raster plots demonstrating cluster activity in the visual neocortex at P9–10. *Insets*: GCaMP3 fluorescence overlaid with binary area plots of three spatially confined cluster events (scale bars, 200 μm). (F–H) Mean CaT frequency per ROI (F), mean cluster area (G), and cluster frequency (H). (I) *Top*: representative traces from a P4 WT of an epoch containing two V1 spindle bursts visible up to 300 μm below the cortical surface. *Bottom*: normalized (mean-subtracted per animal) average whitened power spectra for all V1 spindle bursts recorded at P4 (left) and P7 (right). (J) Occurrence frequencies of V1 spindle bursts were similar between WT and $KO^{Emx1}$ animals at P4 and P7. (K) The average duration of V1 spindle bursts was not significantly different between WT and $KO^{Emx1}$ mice at P4 or P7. (D–K) Each open symbol represents a single animal. Data are presented as mean ± SEM. ns – not significant. $^{*}$ $P < 0.05$, $^{***}$ $P < 0.001$. A synopsis of the applied statistical procedures can be found in Table S6.

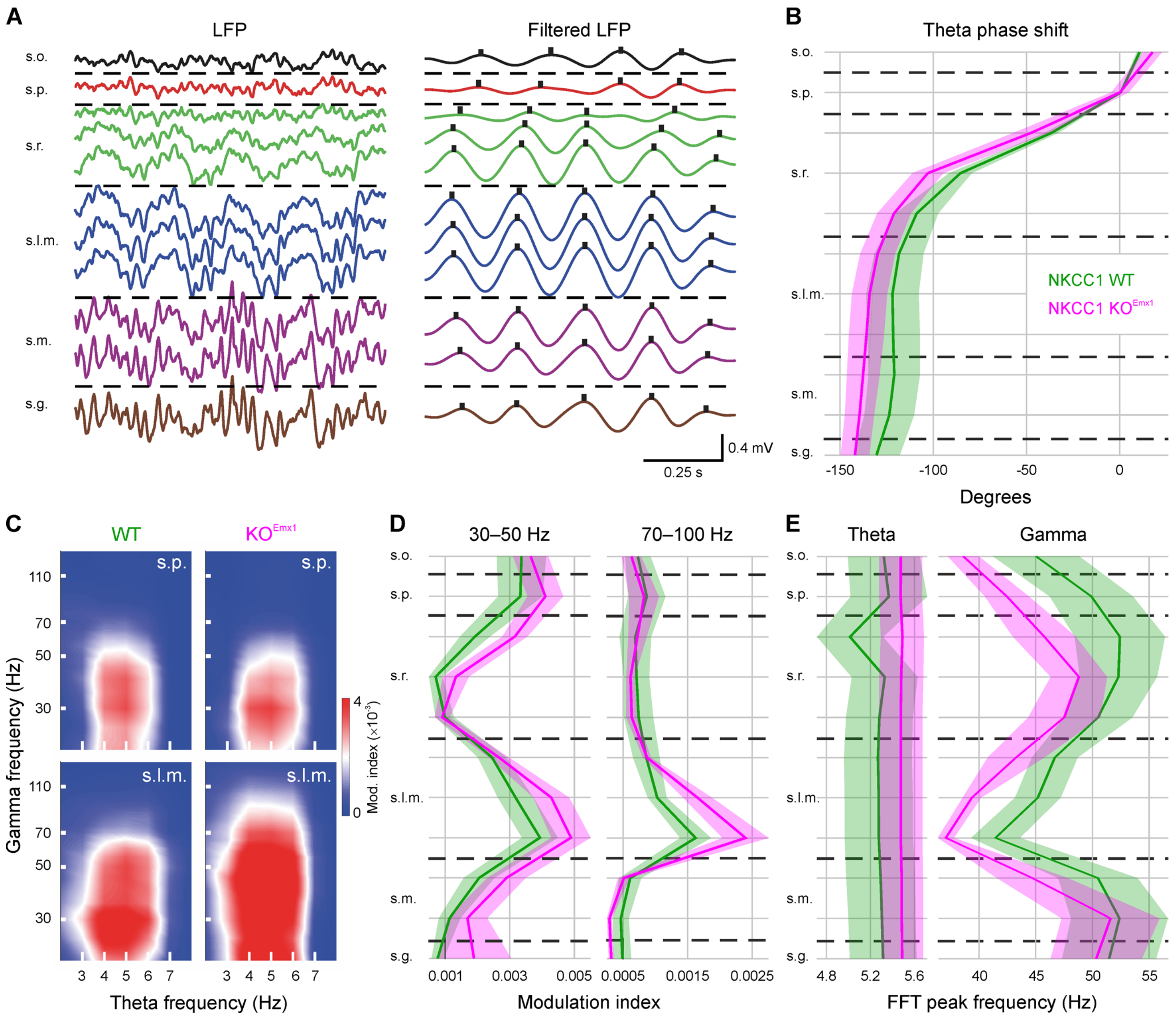


**Figure S7. Related to Figure 7.** Gamma-theta coupling and peak spectral frequencies under urethane anesthesia are unaltered in NKCC1 KO$^{Emx1}$mice. (A) Illustrative example of the theta phase shift across the dorsal hippocampal strata. Right and left panels show the raw and filtered (theta, 3–7 Hz) LFP during a 1s window of REM-like activity, respectively. Note that the peaks of the theta cycles (black dots) show a gradual temporal shift from the s.p. to s.l.m. (B) Similar theta phase shift across the hippocampal strata in WT and KO$^{Emx1}$ mice. Shaded areas represent ± SEM across animals. The s.p. channel was used as a reference. (C) Representative phase-amplitude comodulograms illustrating gamma-theta coupling in s.p. (top) and s.l.m (bottom). (D) Phase-amplitude coupling analysis across the dorsal hippocampal strata. KO$^{Emx1}$ mice showed a trend toward stronger coupling in the high-gamma range (70–100 Hz). (E) Theta and gamma oscillations showed no significant differences in peak spectral frequencies between genotypes. FFT, fast Fourier transform. A synopsis of the applied statistical procedures can be found in Table S7.

**Table S1.** Synopsis of statistical tests related to Figures 1 and S1.

| # | Related to | Descriptive statistics | Test(s) | Test statistics |
|---|---|---|---|---|
| 1 | Figure 1C | WT: 100.0 ± 2.4%, n = 12<br>KO: 38.6 ± 2.4%, n = 13<br>(geometric mean ± SEM) | Mann-Whitney U-test (exact) | $P = 3.8\times10^{-7}$<br>U = 156 |
| 2 | Figure 1E left | WT: 45.0 ± 2.9 m, n = 13<br>KO: 45.2 ± 2.6 m, n = 13 | two-sample t-test | P = 0.94<br>t = –0.075<br>df = 24 |
| 3 | Figure 1E right | WT: 11.5 ± 2.1%, n = 13<br>KO: 9.8 ± 1.1%, n = 13 | Welch's t-test | P = 0.47<br>t = –0.734<br>df = 17.7 |
| 4 | Figure 1G | WT: 68.3 ± 2.7%, n = 15<br>KO: 68.8 ± 2.4%, n = 15 | two-sample t-test | P = 0.89<br>t = –0.145<br>df = 28 |
| 5 | Figure 1I | WT: 213 ± 31 cm, n = 12<br>KO: 244 ± 42 cm, n = 13 | Mann-Whitney U-test (exact) | P = 0.69<br>U = 70 |
| 6 | Figure 1J | WT: n = 12<br>KO: n = 13 | mixed-model ANOVA | interaction: P = 0.31,<br>F = 1.20, df = 7;<br>main effect time: P < 0.0001,<br>F = 22.9, df = 7;<br>main effect genotype: P = 0.96,<br>F = 0.002, df = 1 |
| 7 | Figure 1K | WT: 43.2 ± 3.5%, n = 12<br>KO: 33.1 ± 4.7%, n = 13 | two-sample t-test | P = 0.10<br>t = 1.7<br>df = 23 |
| 8 | Figure 1L | WT: n = 13<br>KO: n = 13 | mixed-model ANOVA | interaction: P = 0.99,<br>F = 0.04, df = 3;<br>main effect time: P < 0.0001,<br>F = 16.46, df = 3;<br>main effect genotype: P = 0.96,<br>F = 1.5, df = 1 |
| 9 | Figure 1M | WT: n = 13<br>KO: n = 13 | mixed-model ANOVA | interaction: P = 0.27,<br>F = 1.3, df = 6;<br>main effect time: P < 0.0001,<br>F = 39.8, df = 6;<br>main effect genotype: P = 0.091,<br>F = 3.1, df = 1 |
| 10 | Figure 1N | WT: n = 13<br>KO: n = 13 | mixed-model ANOVA | interaction: P = 0.43,<br>F = 0.85, df = 2;<br>main effect time: P = 0.065,<br>F = 2.8, df = 2;<br>main effect genotype: P = 0.064,<br>F = 3.5, df = 1 |

**Continuation of Table S1.** Synopsis of statistical tests related to Figures 1 and S1.

| # | Related to | Descriptive statistics | Test(s) | Test statistics |
|---|---|---|---|---|
| 11 | Figure S1K | WT: n = 5-29 (per age-group)<br>KO: n = 6-28 (per age-group) | two-way ANOVA | interaction: $P = 0.61$,<br>$F = 0.77$, df = 7;<br>main effect age: $P = 0$,<br>$F = 410$, df = 7;<br>main effect genotype: $P = 0.35$,<br>$F = 0.88$, df = 1 |
| 12 | Figure S1L | P4: WT: 14.6 ± 3.9 s, n = 6<br>KO: 18.0 ± 2.7 s, n = 9<br>P7: WT: 12.7 ± 4.0 s, n = 8<br>KO: 11.5 ± 3.0 s, n = 7<br>P12: WT: 0.8 ± 0.1 s, n = 6<br>KO: 1.1 ± 0.1 s, n = 8 | two-way ANOVA | interaction: $P = 0.72$,<br>$F = 0.33$, df = 2;<br>main effect age: $P = 1.7 \times 10^{-5}$,<br>$F = 14.8$, df = 2;<br>main effect genotype: $P = 0.72$,<br>$F = 0.13$, df = 1 |
| 13 | Figure S1M | P4: WT: 19.5 ± 4.9 s, n = 6<br>KO: 28.0 ± 1.0 s, n = 9<br>P7: WT: 17.0 ± 3.2 s, n = 8<br>KO: 22.0 ± 4.0 s, n = 7<br>P12: WT: 13.3 ± 3.9 s, n = 6<br>KO: 8.5 ± 1.5 s, n = 8 | two-way ANOVA | interaction: $P = 0.10$,<br>$F = 2.4$, df = 2;<br>main effect age: $P = 8.0 \times 10^{-4}$,<br>$F = 8.7$, df = 2;<br>main effect genotype: $P = 0.26$,<br>$F = 1.3$, df = 1 |
| 14 | lamda-bregma distance | P4 WT: 2.175 ± 0.070 mm, n = 6<br>KO: 2.286 ± 0.135 mm, n = 7<br>P7: WT: 3.063 ± 0.042 mm, n = 8<br>KO: 3.067 ± 0.055 mm, n = 6<br>P12: WT: 3.742 ± 0.058 mm, n = 6<br>KO: 3.644 ± 0.074 mm, n = 8 | two-way ANOVA | interaction: $P = 0.443$,<br>$F = 0.8$, df = 2;<br>main effect age: $P < 0.001$,<br>$F = 164.6$, df = 2;<br>main effect genotype: $P = 0.932$,<br>$F = 0$, df = 1 |

**Table S2.** Synopsis of statistical tests related to Figures 2 and S2.

| # | Related to | Descriptive statistics | Test(s) | Test statistics |
|---|---|---|---|---|
| 1 | Figure 2B<br>Vrest | WT: –73.0 ± 0.6 mV, n = 15<br>KO: –73.3 ± 0.9 mV, n = 10 | Mann-Whitney U-test (exact) | P = 0.77<br>U = 80.5 |
| 2 | Figure 2B<br>Vpeak | WT: –50.9 ± 1.9 mV, n = 15<br>KO: –58.6 ± 2.8 mV, n = 10 | two-sample t-test | P = 0.028<br>t = 2.34<br>df = 23 |
| 3 | Figure 2C | WT: n = 15<br>KO: n = 10 | Spearman's rho | WT: ρ = 0.775, P = $6.9 \times 10^{-4}$<br>KO: ρ = 0.733, P = 0.016 |
| 4 | Figure 2E | WT: baseline: 0.14 ± 0.07, n = 22<br>Isoguvacine: 1.95 ± 0.38, n = 22<br>KO: baseline: 0.19 ± 0.10, n = 16<br>Isoguvacine: 0.31 ± 0.15, n = 16 | paired t-test<br><br>Wilcoxon signed-rank test (exact) | P = $2.6 \times 10^{-4}$, t = -4.39, df = 21<br><br>P = 0.75, W = 5 |
| 5 | Figure 2F<br>Trial 1 | Trial 1: WT: 17 out of 22<br>KO: 3 out of 16 | chi-squared test | P = $3.6 \times 10^{-4}$, $\chi^2$ = 12.7, df = 1 |
| 6 | Figure 2F<br>Trial 2 | Trial 2: WT: 15 out of 22<br>KO: 1 out of 16 | chi-squared test | P = $1.3 \times 10^{-4}$, $\chi^2$ = 14.6, df = 1 |
| 7 | Figure S2E<br>GABA | WT: 23.53 ± 2.05% $\Delta F/F_0$, n = 7<br>KO: 9.10 ± 1.19% $\Delta F/F_0$, n = 6 | two-sample t-test | P = $1.2 \times 10^{-4}$<br>t = 5.82<br>df = 11 |
| 8 | Figure S2E<br>Glutamate | WT: 60.32 ± 6.27% $\Delta F/F_0$, n = 7<br>KO: 57.82 ± 5.07% $\Delta F/F_0$, n = 6 | Mann-Whitney U-test (exact) | P = 0.63<br>U = 25 |
| 9 | Figure S2F | WT: 0.401 ± 0.024, n = 7<br>KO: 0.154 ± 0.014, n = 6 | two-sample t-test | P = $4.1 \times 10^{-6}$<br>t = 8.4<br>df = 11 |

**Table S3.** Synopsis of statistical tests related to Figures 3 and S3.

| # | Related to | Descriptive statistics | Test(s) | Test statistics |
|---|---|---|---|---|
| 1 | Figure 3B | WT: 1.98 ± 0.83 Hz, n = 7<br>KO: 4.40 ± 1.28 Hz, n = 13 | Mann-Whitney U-test (exact) | P = 0.21<br>U = 29 |
| 2 | Figure 3D | WT: 0.34 ± 0.04, n = 6<br>KO: 0.20 ± 0.04, n = 13 | ANCOVA | frequency as a covariate<br>interaction: P = 0.85,<br>F = 0.039, df = 1<br>main effect genotype: P < $1.2\times10^{-4}$,<br>F = 26.3, df = 1;<br>main effect frequency: P = $7.1\times10^{-5}$,<br>F = 29.4, df = 1 |
| 3 | Figure 3F<br>DG | WT: 8.2 ± 1.3%, n = 9<br>KO: 5.3 ± 1.4%, n = 9 | two-sample t-test | P = 0.16<br>t = 1.47<br>df = 16 |
| 4 | Figure 3F<br>CA3 | WT: 11.2 ± 1.9%, n = 9<br>KO: 4.1 ± 1.2%, n = 9 | two-sample t-test | P = $6.4\times10^{-3}$<br>t = 3.133<br>df = 16 |
| 5 | Figure 3F<br>CA1 | WT: 7.2 ± 1.1%, n = 9<br>KO: 4.1 ± 1.2%, n = 9 | Mann-Whitney U-test (exact) | P = 0.064<br>U = 62 |
| 6 | Figure 3F<br>Sub | WT: 8.1 ± 0.9%, n = 9<br>KO: 5.4 ± 2.2%, n = 9 | Mann-Whitney U-test (exact) | P = 0.19<br>U = 56 |
| 7 | Figure 3F<br>Ps | WT: 10.5 ± 1.6%, n = 9<br>KO: 8.3 ± 2.0%, n = 9 | two-sample t-test | P = 0.38<br>t = 0.89<br>df = 16 |
| 8 | Figure 3F<br>MEC | WT: 7.6 ± 1.5%, n = 9<br>KO: 8.0 ± 2.7%, n = 9 | two-sample t-test | P = 0.90<br>t = –0.13<br>df = 16 |
| 9 | Figure 3G<br>DG | WT: 265.4 ± 39.0 $\Delta F/F_0$ × ms, n = 9<br>KO: 50.2 ± 6.7 $\Delta F/F_0$ × ms, n = 7 | Welch's t-test | P = $5.1\times10^{-4}$<br>t = 5.44<br>df = 8.47 |
| 10 | Figure 3G<br>CA3 | WT: 502.8 ± 80.8 $\Delta F/F_0$ × ms, n = 9<br>KO: 50.5 ± 7.6 $\Delta F/F_0$ × ms, n = 7 | Welch's t-test | P = $5.0\times10^{-4}$<br>t = 5.57<br>df = 8.14 |
| 11 | Figure 3G<br>CA1 | WT: 553.7 ± 318.1 $\Delta F/F_0$ × ms, n = 9<br>KO: 50.1 ± 10.6 $\Delta F/F_0$ × ms, n = 5 | Mann-Whitney U-test (exact) | P = $7.7 \times10^{-3}$<br>U = 43 |
| 12 | Figure 3G<br>Sub | WT: 532.8 ± 120.4 $\Delta F/F_0$ × ms, n = 9<br>KO: 39.5 ± 9.2 $\Delta F/F_0$ × ms, n = 6 | Welch's t-test | P = $3.4\times10^{-3}$<br>t = 4.08<br>df = 8.09 |
| 13 | Figure 3G<br>Ps | WT: 1360.4 ± 309.8 $\Delta F/F_0$ × ms, n = 8<br>KO: 115.1 ± 21.3 $\Delta F/F_0$ × ms, n = 8 | Mann-Whitney U-test (exact) | P = $1.4\times10^{-3}$<br>U = 63 |
| 14 | Figure 3G<br>MEC | WT: 1170.2 ± 287.8 $\Delta F/F_0$ × ms, n = 8<br>KO: 90.3 ± 28.5 $\Delta F/F_0$ × ms, n = 8 | Welch's t-test | P = $7.1\times10^{-3}$<br>t = 3.73<br>df = 7.14 |

**Continuation of Table S3.** Synopsis of statistical tests related to Figures 3 and S3.

| # | Related to | Descriptive statistics | Test(s) | Test statistics |
|---|---|---|---|---|
| 15 | Figure S3D | WT: 82.9 ± 6.0%, n = 15<br>KO: 27.0 ± 1.5%, n = 12 | Mann-Whitney U-test | $P = 1.9\times10^{-4}$<br>U = 49.3<br>Z = 3.7 |
| 16 | Figure S3F | WT: 283.7 ± 16.7 ms, n = 8<br>KO: 27.0 ± 1.5 ms, n = 7 | two-sample t-test | $P = 3.0\times10^{-3}$<br>t = –3.64<br>df = 13 |
| 17 | Figure S3G | WT: control: 2.60 ± 0.42 $min^{-1}$, n = 6<br>bumetanide: 0.50 ± 0.12 $min^{-1}$<br>KO: control: 0.83 ± 0.15 $min^{-1}$, n = 8<br>bumetanide: 0.56 ± 0.10 $min^{-1}$ | Welch's t-test (WT control vs KO control)<br>paired t-test (WT)<br>paired t-test (KO) | WT control vs KO control:<br>$P = 6.3\times10^{-3}$, t = 4.03, df = 6.24;<br>WT (cont vs bum): P = 0.0092,<br>t = 4.17, df = 5;<br>KO (cont vs bum): P = 0.015,<br>t = 3.22, df = 7; |
| 18 | Figure S3H | WT: control: 2.46 ± 0.52 $min^{-1}$, n = 6<br>bumetanide: 0.02 ± 0.02 $min^{-1}$<br>KO: control: 0.21 ± 0.10 $min^{-1}$, n = 8<br>bumetanide: 0.04 ± 0.03 $min^{-1}$ | Mann-Whitney U-test (exact) (WT control vs KO control)<br>paired t-test (WT)<br>Wilcoxon signed-rank test (exact, KO) | WT control vs KO control:<br>$P = 6.7\times10^{-4}$, U = 0<br>WT (cont vs bum): P = 0.0047,<br>t = 4.85, df = 5;<br>KO (cont vs bum): P = 0.063,<br>W = 0 |
| 19 | Figure S3I | WT: control: 1.90 ± 0.24 $min^{-1}$, n = 10<br>bumetanide: 0.82 ± 0.17 $min^{-1}$, n = 10<br>KO: control: 1.34 ± 0.23 $min^{-1}$, n = 9<br>bumetanide: 1.17 ± 0.20 $min^{-1}$, n = 9 | mixed-model ANOVA<br>paired t-test (WT)<br>paired t-test (KO) | interaction: P = 0.002,<br>F = 13.86, df = 1;<br>WT bumetanide: $P = 8.1\times10^{-4}$,<br>t = 4.93, df = 9;<br>KO bumetanide: P = 0.058,<br>t = 2.21, df = 8 |
| 20 | Figure S3J | WT: control: 2.14 ± 0.37 $min^{-1}$, n = 10<br>bumetanide: 0.54 ± 0.26 $min^{-1}$, n = 10<br>KO: control: 1.61 ± 0.38 $min^{-1}$, n = 9<br>bumetanide: 1.23 ± 0.25 $min^{-1}$, n = 9 | mixed-model ANOVA<br>paired t-test (WT)<br>Wilcoxon signed-ranked test (exact, KO) | interaction: P = 0.007,<br>F = 9.39, df = 1;<br>WT bumetanide: P = 0.0012,<br>t = 4.63, df = 9;<br>KO bumetanide: P = 0.030,<br>W = 34, Z = 2.17 |

**Table S4.** Synopsis of statistical tests related to Figure 4.

| # | Related to | Descriptive statistics | Test(s) | Test statistics |
|---|---|---|---|---|
| 1 | Figure 4B | WT: 0.315 ± 0.088 Hz, n = 8<br>KO: 0.286 ± 0.113 Hz, n = 11 | Mann-Whitney U-test (exact) | P = 0.39<br>U = 33 |
| 2 | Figure 4D | WT: n = 19 cells<br>KO: n = 16 cells | mixed-model ANOVA | interaction: P = 0.39,<br>F = 1.01, df = 2.59;<br>main effect current injection:<br>$P = 4.0\times10^{-13}$, F = 32.9, df = 2.59;<br>main effect genotype: P = 0.67,<br>F = 0.18, df = 1 |
| 3 | Figure 4E | WT: -44.8 ± 0.7 mV, n = 19<br>KO: -43.6 ± 1.8 mV, n = 16 | Mann-Whitney U-test (exact) | P = 0.8<br>U = 160 |
| 4 | Figure 4G<br>mEPSC frequency | P3–4: WT: 0.30 ± 0.12 Hz, n = 11<br>KO: 0.36 ± 0.08 Hz, n = 13<br>P14-16: WT: 4.7 ± 1.6 Hz, n = 14<br>KO: 3.8 ± 1.0 Hz, n = 27 | two-way ANOVA | interaction: P = 0.69,<br>F = 0.16, df = 1;<br>main effect age: $P = 9.6\times10^{-4}$,<br>F = 12.0, df = 1;<br>main effect genotype: P = 0.73,<br>F = 0.12, df = 1 |
| 5 | Figure 4G<br>mEPSC amplitude | P3–4: WT: 18.0 ± 1.8 pA, n = 11<br>KO: 20.5 ± 2.4 pA, n = 13<br>P14-16: WT: 25.4 ± 2.1 pA, n = 14<br>KO: 27.5 ± 2.1 pA, n = 27 | two-way ANOVA | interaction: P = 0.95,<br>F = 0.004, df = 1;<br>main effect age: $P = 4.3\times10^{-3}$,<br>F = 8.8, df = 1;<br>main effect genotype: P = 0.35,<br>F = 0.90, df = 1 |
| 6 | Figure 4G<br>mEPSC decay | P3–4: WT: 3.8 ± 0.5 ms, n = 11<br>KO: 4.5 ± 0.4 ms, n = 13<br>P14-16: WT: 7.0 ± 0.5 ms, n = 14<br>KO: 7.3 ± 0.4 ms, n = 27 | two-way ANOVA | interaction: P = 0.63,<br>F = 0.24, df = 1;<br>main effect age: $P = 4.8\times10^{-8}$,<br>F = 38.8, df = 1;<br>main effect genotype: P = 0.34,<br>F = 0.93, df = 1 |
| 7 | Figure 4I<br>mIPSC frequency | P3–4: WT: 1.12 ± 0.19 Hz, n = 29<br>KO: 0.62 ± 0.11 Hz, n = 12<br>P14-16: WT: 14.5 ± 0.9 Hz, n = 24<br>KO: 16.1 ± 1.2 Hz, n = 26 | two-way ANOVA | interaction: P = 0.26,<br>F = 1.30, df = 1;<br>main effect age: $P = 1.2\times10^{-27}$,<br>F = 256, df = 1;<br>main effect genotype: P = 0.56,<br>F = 0.35, df = 1 |
| 8 | Figure 4I<br>mIPSC amplitude | P3–4: WT: 57.0 ± 5.4 pA, n = 29<br>KO: 48.4 ± 3.5 pA, n = 12<br>P14-16: WT: 43.0 ± 2.2 pA, n = 24<br>KO: 49.5 ± 2.8 pA, n = 26 | two-way ANOVA | interaction: P = 0.08,<br>F = 3.1, df = 1;<br>main effect age: P = 0.14,<br>F = 2.2, df = 1;<br>main effect genotype: P = 0.82,<br>F = 0.05, df = 1 |
| 9 | Figure 4I<br>mIPSC decay | P3–4: WT: 13.4 ± 0.4 ms, n = 29<br>KO: 14.5 ± 0.7 ms, n = 12<br>P14-16: WT: 9.1 ± 0.5 ms, n = 24<br>KO: 8.4 ± 0.4 ms, n = 26 | two-way ANOVA | interaction: P = 0.07,<br>F = 3.3, df = 1;<br>main effect age: $P = 4.3\times10^{-17}$,<br>F = 110, df = 1;<br>main effect genotype: P = 0.68,<br>F = 0.18, df = 1 |
| 10 | Membrane capacitance | WT: 45.3 ± 3.3 pF, n = 19<br>KO: 55.2 ± 5.2 pF, n = 16 | Mann-Whitney U-test | P = 0.12<br>U = 199<br>Z = 1.54 |
| 11 | Membrane resistance | WT: 0.733 ± 0.067 GΩ, n = 19<br>KO: 0.681 ± 0.075 GΩ, n = 16 | two-sample t-test | P = 0.61<br>t = –0.518<br>df = 33 |
| 12 | Membrane time constant | WT: 33.8 ± 4.4 ms, n = 19<br>KO: 35.5 ± 3.7 ms, n = 16 | two-sample t-test | P = 0.78<br>t = 0.285<br>df = 33 |

**Table S5.** Synopsis of statistical tests related to Figures 5 and S4.

| # | Related to | Descriptive statistics | Test(s) | Test statistics |
|---|---|---|---|---|
| 1 | Figure 5D, Figure S4<br>rad/lm SPW occurrence frequency | P4 WT: 0.1059 ± 0.0125 Hz, n = 6<br>P4 KO: 0.1176 ± 0.0198 Hz, n = 7<br>P7 WT: 0.9274 ± 0.0747 Hz, n = 6<br>P7 KO: 0.8603 ± 0.1001 Hz, n = 5<br>P12 WT: 1.9655 ± 0.1161 Hz, n = 5<br>P12 KO: 1.9869 ± 0.0229 Hz, n = 8 | two-way ANOVA | interaction: $P = 0.741$, $F = 0.3$, df = 2;<br>main effect age: $P < 0.0001$, $F = 496.7$, df = 2;<br>main effect genotype: $P = 0.821$, $F = 0.1$, df = 1 |
| 2 | Figure 5D, Figure S4<br>oriens SPW occurrence frequency | P4 WT: 0.0029 ± 0.0006 Hz, n = 6<br>P4 KO: 0.0034 ± 0.0006 Hz, n = 7<br>P7 WT: 0.0180 ± 0.0050 Hz, n = 6<br>P7 KO: 0.0073 ± 0.0020 Hz, n = 5<br>P12 WT: 0.1599 ± 0.0062 Hz, n = 5<br>P12 KO: 0.2158 ± 0.0254 Hz, n = 8 | two-way ANOVA | interaction: $P = 0.058$, $F = 3.1$, df = 2;<br>main effect age: $P < 0.0001$, $F = 108.5$, df = 2;<br>main effect genotype: $P = 0.201$, $F = 1.7$, df = 1 |
| 3 | Figure 5D, Figure S4<br>rad/lm SPW amplitude | P4 WT: -0.4739 ± 0.0238 mV, n = 6<br>P4 KO: -0.4776 ± 0.0376 mV, n = 7<br>P7 WT: -0.2617 ± 0.0156 mV, n = 6<br>P7 KO: -0.2761 ± 0.0343 mV, n = 5<br>P12 WT: -0.2874 ± 0.0190 mV, n = 5<br>P12 KO: -0.3135 ± 0.0106 mV, n = 8 | two-way ANOVA | interaction: $P = 0.904$, $F = 0.1$, df = 2;<br>main effect age: $P < 0.0001$, $F = 39.24$, df = 2;<br>main effect genotype: $P = 0.484$, $F = 0.5$, df = 1 |
| 4 | Figure 5D, Figure S4<br>oriens SPW amplitude | P4 WT: -0.1886 ± 0.0450 mV, n = 6<br>P4 KO: -0.1529 ± 0.0130 mV, n = 7<br>P7 WT: -0.1764 ± 0.0115 mV, n = 6<br>P7 KO: -0.1455 ± 0.0132 mV, n = 5<br>P12 WT: -0.1292 ± 0.0020 mV, n = 5<br>P12 KO: -0.1367 ± 0.0026 mV, n = 8 | two-way ANOVA | interaction: $P = 0.502$, $F = 0.704$, df = 2;<br>main effect age: $P = 0.158$, $F = 1.961$, df = 2;<br>main effect genotype: $P = 0.242$, $F = 1.424$, df = 1 |
| 5 | Figure 5D<br>HNO occurrence frequency | WT: 0.0457 ± 0.0068 Hz, n = 6<br>KO: 0.0570 ± 0.0065 Hz, n = 7 | one-way ANOVA | genotype: $P = 0.252$, $F = 1.46$, df = 1 |
| 6 | Figure 5D<br>HNO mean length | WT: 1.9287 ± 0.2206 s, n = 6<br>KO: 2.2805 ± 0.1159 s, n = 7 | one-way ANOVA | genotype: $P = 0.169$, $F = 2.173$, df = 1 |

**Table S6.** Synopsis of statistical tests related to Figures 6, S5 and S6.

| # | Related to | Descriptive statistics | Test(s) | Test statistics |
|---|---|---|---|---|
| 1 | Figure 6F | WT: 5.81 ± 0.30 $min^{-1}$, n = 12<br>KO: 4.56 ± 0.37 $min^{-1}$, n = 10 | two-sample t-test | P = 0.015<br>t = 2.66<br>df = 20 |
| 2 | Figure 6G | +SPW: WT: 2.05 ± 0.14 $min^{-1}$, n = 12<br>KO: 1.68 ± 0.25 $min^{-1}$, n = 10<br>–SPW: WT: 5.82 ± 0.36 $min^{-1}$, n = 12<br>KO: 4.62 ± 0.33 $min^{-1}$, n = 10 | mixed-model ANOVA<br><br><br><br><br>two-sample t-test<br>two-sample t-test | interaction: P = 0.14,<br>F = 2.4, df = 1;<br>main effect SPW: P = 5.7×$10^{-11}$,<br>F = 159, df = 1;<br>main effect genotype: P = 0.018,<br>F = 6.7, df = 1<br>P (+SPW WT vs KO) = 0.19<br>P (–SPW WT vs KO) = 0.027 |
| 3 | Figure 6H | +SPW: WT: 3.45 ± 0.25 $\Delta F/F_0$ × $10^{-3}$ × $SI^{-1}$<br>KO: 3.84 ± 0.34 $\Delta F/F_0$ × $10^{-3}$ × $SI^{-1}$<br>–SPW: WT: 1.67 ± 0.37 $\Delta F/F_0$ × $10^{-3}$ × $SI^{-1}$<br>KO: 1.80 ± 0.74 $\Delta F/F_0$ × $10^{-3}$ × $SI^{-1}$ | mixed-model ANOVA | interaction: P = 0.45,<br>F = 0.60, df = 1;<br>main effect SPW: P = 6.6×$10^{-10}$,<br>F = 120, df = 1;<br>main effect genotype: P = 0.28,<br>F = 1.22, df = 1 |
| 4 | Figure 6I | +SPW: WT: 91.1 ± 2.2%<br>KO: 88.7 ± 2.9%<br>–SPW: 59.0 ± 1.3%<br>KO: 58.8 ± 2.5% | mixed-model ANOVA | interaction: P = 0.48,<br>F = 0.52, df = 1;<br>main effect SPW: P = 7.3×$10^{-15}$,<br>F = 416, df = 1;<br>main effect genotype: P = 0.67,<br>F = 0.19, df = 1 |
| 5 | Figure 6J | +SPW: WT: 3.73 ± 0.21% $\Delta F/F_0$<br>KO: 3.79 ± 0.38% $\Delta F/F_0$<br>–SPW: WT: 2.64 ± 0.06% $\Delta F/F_0$<br>KO: 2.68 ± 0.08% $\Delta F/F_0$ | mixed-model ANOVA | interaction: P = 0.96,<br>F = 0.0024, df = 1;<br>main effect SPW: P = 4.0×$10^{-6}$,<br>F = 39.3, df = 1;<br>main effect genotype: P = 0.84,<br>F = 0.04, df = 1 |
| 6 | Figure 6L | +SPW: control: 1.72 ± 0.27 $min^{-1}$, n = 6<br>gabazine: 12.40 ± 1.11 $min^{-1}$,<br>–SPW: control: 5.56 ± 0.68 $min^{-1}$, n = 6<br>gabazine: 3.15 ± 0.85 $min^{-1}$, | two-way repeated-measures ANOVA<br><br><br><br>paired t-test<br>paired t-test | interaction: P = 6.2×$10^{-4}$,<br>F = 57.9, df = 1;<br>main effect SPW: P = 0.055,<br>F = 6.2, df = 1;<br>main effect gabazine: P = 3.4×$10^{-4}$,<br>F = 75.0, df = 1<br>P (+SPW cont vs Gbz) = 2.2×$10^{-4}$<br>P (–SPW cont vs Gbz) = 0.031 |
| 7 | Results section (related to Figure 6L) Amplitude of Ca2+ clusters | +SPW: control: 3.74 ± 0.26% $\Delta F/F_0$<br>gabazine: 40.1 ± 8.3% $\Delta F/F_0$<br>–SPW: control: 3.1 ± 0.21% $\Delta F/F_0$<br>gabazine: 11.5 ± 3.6% $\Delta F/F_0$ | two-way repeated-measures ANOVA<br><br><br><br>paired t-test<br>Wilcoxon signed-ranked test | interaction: P = 0.038,<br>F = 7.85, df = 1;<br>main effect SPW: P = 0.028,<br>F = 9.3, df = 1;<br>main effect gabazine: P = 3.2×$10^{-3}$,<br>F = 28.1, df = 1<br>P (+SPW cont vs Gbz) = 7.7×$10^{-3}$<br>P (–SPW cont vs Gbz) = 0.036 |
| 8 | Figure 6N | WT: 2.48 ± 0.04 $min^{-1}$, n = 6<br>KO: 2.48 ± 0.14 $min^{-1}$, n = 7 | Welch's t-test | P = 1<br>t = –5.4 × $10^{-4}$<br>df = 7.05 |
| 9 | Figure 6O | WT: 12.8 ± 0.7%, n = 6<br>KO: 11.5 ± 0.7%, n = 7 | two-sample t-test | P = 0.21<br>t = –1.2<br>df = 11 |
| 10 | Figure 6P | WT: 19.7 ± 1.3 $min^{-1}$, n = 6<br>KO: 21.7 ± 1.1 $min^{-1}$, n = 7 | two-sample t-test | P = 0.25<br>t = 1.33<br>df = 11 |

**Continuation of Table S6.** Synopsis of statistical tests related to Figures 6, S5 and S6.

| # | Related to | Descriptive statistics | Test(s) | Test statistics |
|---|---|---|---|---|
| 11 | Figure S5C<br>all SPWs | Transcortical: 2.82 ± 0.37 $min^{-1}$, n = 6<br>Window: 2.42 ± 0.16 $min^{-1}$, n = 4 | Mann-Whitney U-test (exact) | P = 0.91<br>U = 13<br>Z = 0.11 |
| 12 | Figure S5C<br>SPW wo movement | Transcortical:1.12 ± 0.15 $min^{-1}$, n = 6<br>Window: 1.58 ± 0.25 $min^{-1}$, n = 4 | two-sample t-test | P = 0.15<br>t = –1.61<br>df = 8 |
| 13 | Figure S5C<br>SPW w movement | Transcortical: 1.71 ± 0.31 $min^{-1}$, n = 6<br>Window: 0.87 ± 0.17 $min^{-1}$, n = 4 | two-sample t-test | P = 0.075<br>t = 2.05<br>df = 8 |
| 14 | Figure S5E<br>all SPWs | control: 2.99 ± 0.43 $min^{-1}$, n = 4<br>TTX: 0 ± 0 $min^{-1}$ | paired t-test | P = 0.006<br>t = 7.01<br>df = 3 |
| 15 | Figure S5E<br>all CaTs | control: 4.47 ± 0.47 $min^{-1}$, n = 4<br>TTX: 0.01 ± 0.01 $min^{-1}$ | paired t-test | P = $3.0 \times 10^{-4}$<br>t = 19.3<br>df = 3 |
| 16 | Figure S6C<br>Trial 1 | WT: 10 out of 22<br>KO: 2 out of 18 | chi-squared test | P = 0.018, $\chi^2$ = 5.56, df = 1 |
| 17 | Figure S6C<br>Trial 2 | WT: 12 out of 22<br>KO: 3 out of 18 | chi-squared test | P = 0.014, $\chi^2$ = 6.06, df = 1 |
| 18 | Figure S6C<br>Trial 3 | WT: 12 out of 22<br>KO: 3 out of 18 | chi-squared test | P = 0.014, $\chi^2$ = 6.06, df = 1 |
| 19 | Figure S6D | WT: 100.0 ± 2.9%, n = 8<br>KO: 18.9 ± 2.3%, n = 8<br>(geometric mean ± SEM) | Mann-Whitney U-test (exact) | P = $9.4 \times 10^{-4}$<br>U = 64 |
| 20 | Figure S6F | WT: 9.6 ± 0.3 $min^{-1}$, n = 7<br>KO: 8.9 ± 0.2 $min^{-1}$, n = 8 | Mann-Whitney U-test (exact) | P = 0.073<br>U = 44 |
| 21 | Figure S6G | WT: 23.4 ± 1.0%, n = 7<br>KO: 23.4 ± 0.5%, n = 8 | two-sample t-test | P = 0.98<br>t = 0.023<br>df = 13 |
| 22 | Figure S6H | WT: 41.1 ± 2.1 $min^{-1}$, n = 7<br>KO: 37.9 ± 1.4 $min^{-1}$, n = 8 | two-sample t-test | P = 0.21<br>t = 1.31<br>df = 13 |
| 23 | Figure S6J | P4 WT: 0.0186 ± 0.0019 Hz, n = 6<br>P4 KO: 0.0248 ± 0.0023 Hz, n = 7<br>P7 WT: 0.0081 ± 0.0010 Hz, n = 7<br>P7 KO: 0.0091 ± 0.0019 Hz, n = 5 | two-way ANOVA | Interaction P = 0.183<br>F = 1.9, df = 1;<br>main effect genotype: P = 0.067<br>F = 3.73, df = 1 |
| 23 | Figure S6K | P4 WT: 5.81 ± 0.28 s, n = 6<br>P4 KO: 5.74 ± 0.30 s, n = 7<br>P7 WT: 5.54 ± 0.62 s, n = 7<br>P7 KO: 3.80 ± 0.36 s, n = 5 | two-way ANOVA | Interaction P = 0.071<br>F = 3.609, df = 1;<br>main effect genotype: P = 0.053<br>F = 4.21, df = 1 |

**Table S7.** Synopsis of statistical tests related to Figures 7 and S7.

| # | Related to | Descriptive statistics | Test(s) | Test statistics |
|---|---|---|---|---|
| 1 | Figure 7B<br>Band-limited-power Amplitude | theta (3 - 7 Hz) band:<br>WT ch 6 (s.l.m): 0.43 ± 0.05 mV, n = 7<br>KO ch 6 (s.l.m): 0.57 ± 0.05 mV, n = 5<br>WT ch 7 (s.l.m): 0.48 ± 0.06 mV, n = 7<br>KO ch 7 (s.l.m): 0.67 ± 0.06 mV, n = 5<br>WT ch 8 (s.l.m): 0.45 ± 0.05 mV, n = 7<br>KO ch 8 (s.l.m): 0.66 ± 0.06 mV, n = 5<br>WT ch 9 (s.m): 0.40 ± 0.05 mV, n = 7<br>KO ch 9 (s.m.): 0.58 ± 0.05 mV, n = 5<br>WT ch 10 (s.m.): 0.32 ± 0.04 mV, n = 7<br>KO ch 10 (s.m.): 0.48 ± 0.04 mV, n = 5 | nested repeated measures ANOVA<br><br>subgroups: theta/gamma/ MUA -band<br>measures: each HC channel group | interact. band*chan*genotype: P < 0.001, F = 4.2, df = 20;<br>interact. channel*genotype: P = 0.001, F = 3.4, df = 10;<br>interact. band*genotye: P = 0.007, F = 6.5, df = 2;<br>main effect genotype: P = 0.055, F = 4.7, df = 1;<br>Tukey HSD post-hoc, WT vs. KO, theta (3 - 7 Hz) band:<br>ch 6: P = 0.03517<br>ch 7: P = 0.00133<br>ch 8: P = 0.00046<br>ch 9: P = 0.00231<br>ch 10: P = 0.01717 |
| 2 | Figure 7D<br>ripple spectral frequency | WT: 113 ± 1.6 Hz, n = 8<br>KO: 108 ± 1.5 Hz, n = 10 | one-way ANOVA | P = 0.036<br>F = 5.246<br>df = 15 |
| 3 | Figure 7D<br>SPW amplitude | WT: -2.189 ± 0.216 mV, n = 8<br>KO: -1.939 ± 10.288 mV, n = 10 | one-way ANOVA | P = 0.516<br>F = 0.441<br>df = 1 |
| 4 | Figure 7D<br>SPW incidence frequency | WT: 0.247 ± 0.034 Hz, n = 8<br>KO: 0.140 ± 0.022 Hz, n = 10 | one-way ANOVA | P = 0.015<br>F = 7.408<br>df = 1 |
| 5 | Figure 7H | WT: 3.23 ± 0.35 $min^{-1}$, n = 9<br>KO: 3.04 ± 0.33 $min^{-1}$, n = 8 | two-sample t-test | P = 0.69<br>t = 0.41<br>df = 15 |
| 6 | Figure 7I | measured: WT: 0.0077 ± 0.0008, n = 9<br>KO: 0.0055 ± 0.0004, n = 8<br>shuffled: WT: $5.2\times10^{-6}$ ± $2.0\times10^{-6}$, n = 9<br>KO: $5.2\times10^{-6}$ ± $1.8\times10^{-6}$, n = 8 | two-sample t-test (measured) | P = 0.029<br>t = 2.41<br>df = 15 |
| 7 | Figure 7K | measured: WT: 0.0094 ± 0.0010, n = 9<br>KO: 0.0064 ± 0.0006, n = 8<br>shuffled: WT: $3.1\times10^{-5}$ ± $7.9\times10^{-6}$, n = 9<br>KO: $1.4\times10^{-5}$ ± $6.3\times10^{-6}$, n = 8 | two-sample t-test (measured) | P = 0.29<br>t = 2.41<br>df = 15 |
| 8 | Figure S7B<br>Theta phase shift | WT: n = 7<br>KO: n = 5 | repeated measures ANOVA | Interaction P = 0.705, F = 0.7, df = 10;<br>main effect genotype: P = 0.402<br>F = 0.8, df = 1 |
| 9 | Figure S7D<br>Modulation Index | WT: n = 7<br>KO: n = 5 | nested repeated measures ANOVA<br>subgroups: gamma band group | Interaction P = 0.508, F = 0.93, df = 10;<br>main effect genotype: P = 0.315<br>F = 1.12, df = 1 |
| 10 | Figure S7E<br>Theta FFT peak | WT: n = 7<br>KO: n = 5 | repeated measures ANOVA | Interaction P = 0.832, F = 0.573, df = 10;<br>main effect genotype: P = 0.613<br>F = 0.273, df = 1 |
| 11 | Figure S7E<br>Gamma FFT peak | WT: n = 7<br>KO: n = 5 | repeated measures ANOVA | Interaction P = 0.769, F = 0.649, df = 10;<br>main effect genotype: P = 0.274<br>F = 1.339, df = 1 |

**Table S8.** Antibodies, mouse strains and chemicals used in this study.

| Resource | Source | ID |
|---|---|---|
| **Antibodies** | | |
| Mouse anti-NeuN (1:500) | Chemicon | Cat #: MAB377<br>RRID: AB_2298772 |
| Mouse anti-GAD67 (1:2,500) | Millipore | Cat #: MAB5406<br>RRID: AB_2278725 |
| Rabbit anti-GFAP (1:500) | DAKO | Cat #: Z0334<br>RRID: AB_10013382 |
| Rabbit anti-parvalbumin (1:1,000) | Swant | Cat #: PV 25<br>RRID: AB_10000344 |
| Mouse anti-somatostatin (1:100) | Acris | Cat #: DM3200<br>RRID: n.a. |
| Rabbit anti-NeuN (1:500) | Abcam | Cat #: ab104225<br>RRID: AB_10711153 |
| Alexa Fluor 488 donkey anti-mouse (1:500) | Invitrogen | Cat #: A-21202<br>RRID: AB_141607 |
| Alexa Fluor 488 donkey anti-rabbit (1:500) | Invitrogen | Cat #: A-21206<br>RRID: AB_141708 |
| Alexa Fluor 647 donkey anti-rabbit (1:500) | Dianova | Cat #: 715-605-151<br>RRID: AB_2340863 |
| **Mouse strains** | | |
| *Emx1*$^{IREScre}$ (B6.129S2-*Emx1*$^{tm1(cre)Krj}$/J) | The Jackson Laboratory | Stock #: 005628<br>RRID: MGI_3617405 |
| Ai38 (B6;129S6-*Gt(ROSA)26Sor*$^{tm38(CAG-GCaMP3)Hze}$/J) | The Jackson Laboratory | Stock #: 014538<br>RRID: MGI_4999582 |
| Ai14 (B6;129S6-*Gt(ROSA)26Sor*$^{tm14(CAG-tdTomato)Hze}$/J) | The Jackson Laboratory | Stock #: 007908<br>RRID: MGI_3817869 |
| *Slc12a2*$^{flox/flox}$ | C. Hübner, Institute of Human Genetics, University Hospital Jena | Antoine et al.<br>Science 2013 |
| C57BL/6J | The Jackson Laboratory | Stock #: 000664<br>RRID: MGI_3028467 |
| **Chemicals and Substances** | | |
| 1(S),9(R)-(−)-Bicuculline methiodide (BMI) | Sigma-Aldrich | CAS #: 40709-69-1<br>Cat #: 14343 |
| Gabazine (SR-95531) | Sigma-Aldrich | CAS #: 104104-50-9<br>Cat #: S106 |
| Bumetanide | Sigma-Aldrich | CAS #: 28395-03-1<br>Cat #: B3023 |
| DL-2-amino-5-phosphonopentanoic acid (APV) | Tocris | CAS #: 76326-31-3<br>Cat #: 0105 |
| 6,7-dinitroquinoxaline-2,3(1H,4H)-dione (DNQX) | Tocris | CAS #: 2379-57-9<br>Cat #: 0189 |
| Tetrodotoxin citrate (TTX) | Biotrend | CAS #: 18660-81-6<br>Cat #: BN0518 |
| γ-Aminobutyric acid (GABA) | Sigma-Aldrich | CAS #: 56-12-2<br>Cat #: A2129 |
| L-Glutamic acid monosodium salt monohydrate | Sigma-Aldrich | CAS #: 6106-04-3<br>Cat #: 49621 |
| Isoguvacine hydrochloride | Tocris | CAS #: 68547-97-7<br>Cat #: 0235 |
| Oregon Green 488 BAPTA-1, AM (OGB1) | Invitrogen | CAS #: 244167-57-5<br>Cat #: O6807 |
